# Engineering chimeric antigen receptor neutrophils from human pluripotent stem cells for targeted cancer immunotherapy

**DOI:** 10.1101/2022.03.02.482679

**Authors:** Yun Chang, Ramizah Syahirah, Xuepeng Wang, Gyuhyung Jin, Sandra E. Torregrosa-Allen, Bennett D. Elzey, Sydney N Hummel, Tianqi Wang, Xiaojun Lian, Qing Deng, Hal E. Broxmeyer, Xiaoping Bao

## Abstract

Neutrophils, the most abundant white blood cells in the circulation, are closely related to cancer development and progression. Primary neutrophils from healthy donors present potent cytotoxicity against different human cancer cell lines through direct contact and via the generation of reactive oxygen species (ROS). However, due to their short half-life and resistance to genetic modification, neutrophils have not yet been engineered with widely used chimeric antigen receptors (CARs) to enhance their anti-tumor cytotoxicity for targeted immunotherapy. Here, we genetically engineered human pluripotent stem cells (hPSCs) with different synthetic CARs and successfully differentiated them into functional neutrophils by implementing a novel chemically-defined differentiation platform. Neutrophils expressing the chlorotoxin (CLTX)-T-CAR presented specific cytotoxicity against glioblastoma (GBM) cells both in monolayer and 3D cultures. In a GBM xenograft mouse model, systematically-administered CLTX-T-CAR neutrophils also displayed enhanced anti-tumor activity and prolonged animal survival compared with peripheral blood-neutrophils, hPSC-neutrophils and CLTX-NK-CAR natural killer (NK) cells. Collectively, we established a new platform for production of CAR-neutrophils, paving the way to myeloid cell-based therapeutic strategies that would complement and boost current cancer treatment approaches.

## INTRODUCTION

Neutrophils, the most abundant circulating leukocytes in humans, accumulate in many types of tumors and represent a significant portion of tumor-infiltrating cells (Eruslanov et al., 2017; Ilie et al., 2012; Jaillon et al., 2020; Zhao et al., 2020). Due to their heterogeneity and plasticity in the tumor microenvironment (TME), neutrophils have demonstrated contradictory pro-tumor and anti-tumor effects during tumor evolution. For instance, tumor-associated neutrophils (TANs) present direct or antibody-dependent cytotoxicity against solid cancers (Kargl et al., 2019; Matlung et al., 2018), whereas they also facilitate angiogenesis, promote tumor cell migration, and suppress anti-tumor functions of other immune cells in the TME (Coffelt et al., 2015, 2016; Huo et al., 2019). The presence of pro-tumor neutrophils has limited the efficacy of many cancer therapies (Itatani et al., 2020), including emerging immunotherapies, leading to development of suppressive neutrophil-targeted strategies for treating various cancers in both preclinical studies and clinical trials (Zhao et al., 2020). However, given the high heterogeneity of neutrophils in the complex TME (Lecot et al., 2019; Sagiv et al., 2015), general suppression with small molecule- or antibody-based approaches may eliminate both pro-tumor and anti-tumor neutrophils, and reduce efficacy of neutrophil-targeted therapy. Furthermore, adverse effects, such as neutropenia (McDermott et al., 2010), may develop in cancer patients and increase their risk of infections (Xie et al., 2020). Thus, alternative neutrophil-targeting approaches are needed to realize their full potential in cancer treatment.

For the past decade, chimeric antigen receptors (CARs) have been developed and widely used in T and natural killer (NK) cells to boost their anti-tumor effects on various hematologic malignancies and solid tumors (Feins et al., 2019; June and Sadelain, 2018; June et al., 2018; Lim and June, 2017; Mehta and Rezvani, 2018; Zhu et al., 2018), revolutionizing the field of cancer immunotherapy.

More recently, genetic engineering of primary macrophages with CARs has programmed them as anti-tumor effector cells and improved phagocytosis (Klichinsky et al., 2020). Given their similarity to macrophages and shared innate anti-tumor responses (Yan et al., 2014), we hypothesized that human neutrophils may present enhanced tumoricidal activities in various ways after CAR engineering. Indeed, CD4ζ chimeric immune receptors (CIRs) improved the cytolysis of human neutrophils against HIVenv-transfected cells *in vitro* with a lysis efficiency of ∼10% at an effector-to-target (E:T) ratio of 10:1 (Roberts et al., 1998), which is possibly due to gene silencing during neutrophil differentiation from CD34+ hematopoietic progenitors. These prior studies encouraged our interest in the search of alternative cell sources and preparation approaches for potent CAR-neutrophils in treating solid tumors.

To address these challenges, we first developed a chemically-defined, feeder-free platform for robust generation of functional neutrophils from human pluripotent stem cells (hPSCs) by applying stage-specific morphogens. Based on previous studies on CAR designs against solid tumors (Kim et al., 2020a; Li et al., 2018; Nguyen et al., 2012; Wang et al., 2020a), we synthesized and knocked 3 different glioblastoma (GBM)-targeting CAR constructs into the endogenous *AAVS1* safe harbor locus in H9 hPSCs by CRISPR/Cas9-mediated homologous recombination, and evaluated their ability in improving neutrophil-mediated tumor killing. We found that the CLTX-T-CAR construct, composed of chlorotoxin (CLTX) (Wang et al., 2020a), a 36-amino acid GBM-targeting peptide found in *Leiurus quinquestriatus* scorpion venom (Qin et al., 2014), the transmembrane domain of CD4 and the intracellular domain of CD3ζ, as best in enhancing anti-tumor cytotoxicity of hPSC-derived neutrophils. The resulting CLTX-T-CAR neutrophils presented a typical neutrophil phenotype and killed the targeted tumor cells through specific binding to GBM via membrane-associated matrix metalloproteinase 2 (MMP2). Compared with PBS control, peripheral blood (PB) neutrophils, wildtype hPSC-derived neutrophils, wildtype NK and CAR-NK cells, systematically administered CLTX-T-CAR hPSC-derived neutrophils significantly inhibited tumor growth in the *in situ* GBM xenograft models and prolonged animal survival. Collectively, our neutrophil differentiation platform in combination with genome editing techniques may provide a realistic approach to manufacture off-the-shelf CAR-neutrophils for targeted immunotherapy, thus paving the way to myeloid cell-based therapeutic strategies that would complement and enhance the efficacy of cancer immunotherapy.

## RESULTS

### Chemically-defined condition allows robust generation of functional neutrophils

*In vitro* hematopoietic progenitor induction is the first step to generate neutrophils from hPSCs (Brok-Volchanskaya et al., 2019; Lachmann et al., 2015; Saeki et al., 2009; Sweeney et al., 2016; Trump et al., 2019). Multipotent hematopoietic stem and progenitor cells (HSPCs) arise from the arterial vasculature in the aorta-gonad-mesonephros (AGM) region through endothelial-to-hematopoietic transition (EHT) (Bertrand et al., 2010; Boisset et al., 2010; Kissa and Herbomel, 2010). We previously induced homogenous CD34+CD31+ hemogenic endothelium (HE) from hPSCs via small-molecule activation of Wnt signaling alone (**Fig. S1A-S1B**) (Bao et al., 2015; Lian et al., 2014). Importantly, the resulting HE also expressed SOX17 (**Fig. S1B-S1D**), a transcription factor expressed in vascular structures of the AGM and required for HSC generation from AGM (Clarke et al., 2013; Kim et al., 2007; Ng et al., 2016). The employment of TGFβ inhibitor SB431542 (SB) (Bertrand et al., 2010; Boisset et al., 2010; Kissa and Herbomel, 2010) promoted EHT process for the generation of CD45+CD43+ HSPCs (**Fig. S1E-F**) that co-expressed definitive hematopoiesis markers CD44 (Fidanza et al., 2019; Oatley et al., 2020) (**Fig. S1G**) and RUNX1c (Ng et al., 2016) (**Fig. S1H**). The resulting day 15 hPSC-derived HSPCs also maintained high viability after freeze-thaw (**Fig. S1I-J**).

To induce myeloid progenitor and neutrophil differentiation, hPSC-derived HSPC cultures were treated with granulocyte-macrophage colony-stimulating factor (GM-CSF), IL-3, and IL-6 (Cao et al., 2019) from day 9 (**Fig. S2A**). Floating myeloid progenitor cells collected at different days presented both granulocyte-macrophage (GM) and macrophage (M)-colony-forming potential (**Fig. S2B-C**), which increased from day 12 to day 18 and decreased afterwards. To promote neutrophil specification, day 15 myeloid progenitors were treated with granulocyte-CSF (G-CSF). As expected, G-CSF significantly decreased the number of CD14+ monocytes/macrophages as compared to GM-CSF (**Fig. S2D-E**). To identify optimal myeloid progenitors for neutrophil differentiation, floating cells collected at days 12, 15 and 18 were subjected to G-CSF treatment along with AM580 (**Fig. S2F**), a retinoic acid agonist that promotes neutrophil production from human CD34+ cells (Brok-Volchanskaya et al., 2019; Li et al., 2016). The efficiency of neutrophil differentiation increased from day 15 to day 21 and significantly decreased afterward, which may be due to the short life-span of neutrophils (**Fig. S2G-H**). Next, we identified day 15 myeloid progenitors with 6-day treatment of G-CSF and AM580 as the optimal condition for neutrophil differentiation, which was thus employed in our subsequent experiments (**Fig. 1A**). We also assessed the phenotype of sorted CD16-cells (**Fig. S2I**) in the neutrophil differentiation culture, which were mainly composed of ∼20% FcεR1α+ basophils and ∼74% EPX+ eosinophils (Choi et al., 2009) (**Fig. S2J**). Dynamic morphological changes along with the emergence of hematopoietic clusters from day 12 were observed (**Fig. 1B**). The resulting day 21 neutrophils displayed a typical neutrophil morphology (**Fig. S2K-L**) and manifested high expression levels of neutrophil-specific markers (**Fig. 1C**), including CD16, CD11b, CD15, CD66b, CD18 and MPO, as compared to their counterparts isolated from human peripheral blood (PB).

**Fig. 1.**
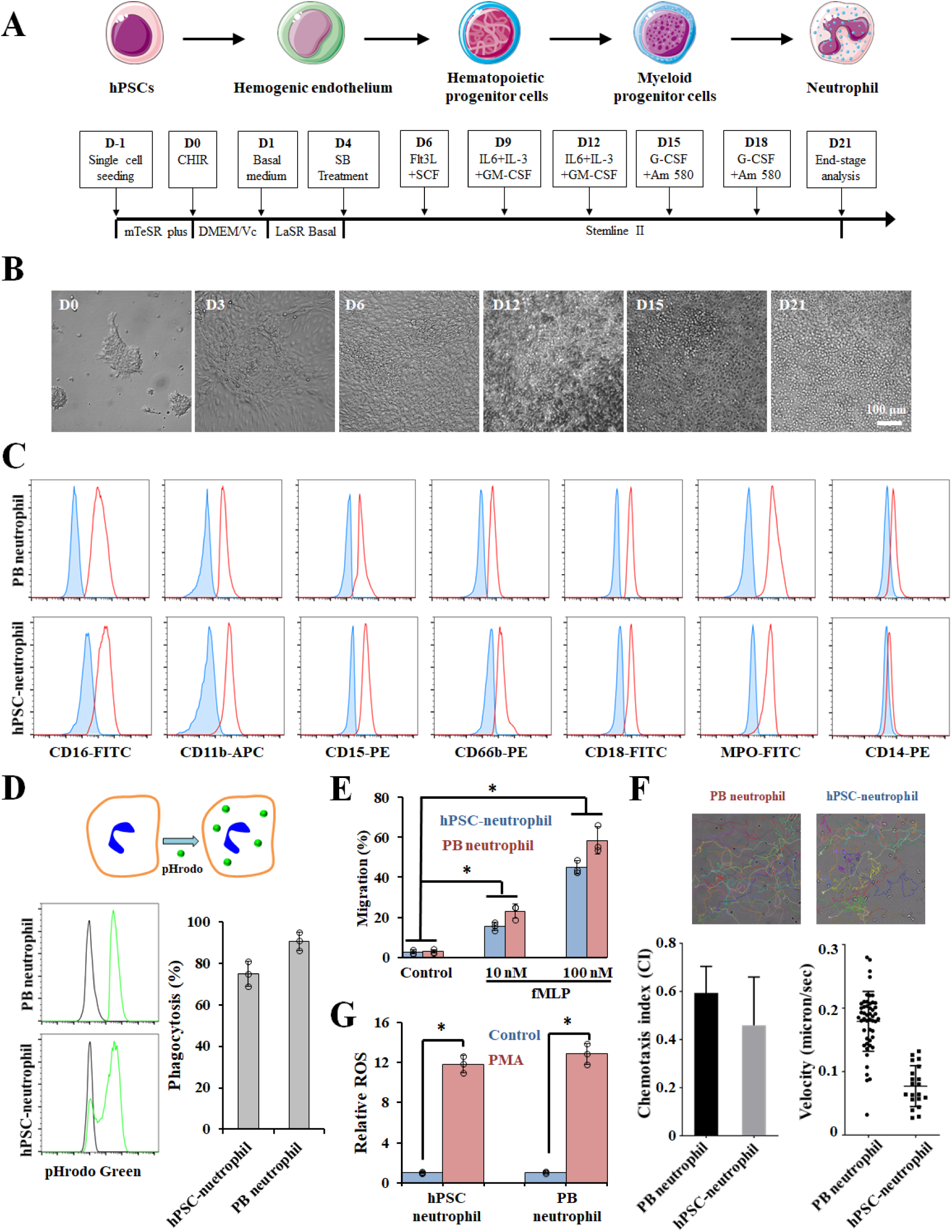
hPSC-derived neutrophils adopt a molecular and functional phenotype similar to primary neutrophils. (A) Schematic of optimized neutrophil differentiation from hPSCs under chemically-defined conditions. (B) Representative bright-field images of cellular morphology at the indicated days: D0 hPSCs, D3 mesoderm, D6 hemogenic endothelium, D12 hematopoietic stem and progenitor cells (HSPCs), D15 myeloid progenitor cells, and D21 neutrophils. Scale bars, 100 μm. (C) Flow cytometric analysis of generated neutrophils. Plots show unstained control (blue) and specific antibody (red) histograms. Primary peripheral blood (PB) neutrophils were used as a positive control. (D) Phagocytosis of pHrodo Green *E. coli* particles by hPSC-derived neutrophils. (E) Transwell migration analysis of the hPSC-derived neutrophils in the absence or presence of chemoattractant (10 nM and 100 nM of fMLP). Data are represented as mean ± s.d. of three independent replicates, \**p*<0.05. (F) Representative tracks, mean velocity, and chemotaxis index of human PB and hPSC-derived neutrophils during chemotaxis are shown. (G) Reactive oxygen species (ROS) production of hPSC-derived and primary PB neutrophils with or without phorbol 12-myristate 13-acetate (PMA) treatment. Data are represented as mean ± s.d. of three independent replicates, \**p*<0.05. CHIR: CHIR99021; SB: SB431542; SCF: stem cell factor; G-CSF: granulocyte colony-stimulating factor; GM-CSF: granulocyte-macrophage colony-stimulating factor; Vc: vitamin C or ascorbic acid.

To evaluate the function of hPSC-derived neutrophils, we performed phagocytosis and chemotaxis assays. Similar to primary PB neutrophils, hPSC-derived neutrophils effectively phagocytosed pHrodo *E. coli* bioparticles (**Fig. 1D**) and displayed excellent migration ability in chemotaxis models using transwells (**Fig. 1E**) and microfluidics (**Fig. 1F**) (Afonso et al., 2013). We also measured the production of reactive oxygen species (ROS) from hPSC-derived neutrophils. In response to the phorbol 12-myristate 13-acetate (PMA), hPSC-derived neutrophils generated comparable ROS to PB neutrophils (**Fig. 1G**). Collectively, we established a chemically-defined, feeder-free platform for robust production of functional neutrophils from hPSCs with a yield of ∼20 neutrophils per hPSC, thus highlighting its potential applications in studying neutrophil biology and treating neutropenia.

### *AAVS1* targeted CAR knockin improves anti-tumor cytotoxicity of hPSC-derived neutrophils

Primary neutrophils from healthy donors present potent cancer-killing activity against various human cancer cell lines (Yan et al., 2014). Thus, we sought to determine whether *de novo* neutrophils derived from hPSCs are able to directly kill tumor cells, and whether chimeric antigen receptor (CAR) expression could enhance their anti-tumor cytotoxicity for targeted immunotherapy. To achieve stable and uniform expression of CARs on neutrophils, we directly knocked CAR constructs into the endogenous *AAVS1* safe harbor locus of H9 hPSCs, a widely-used site for constitutive transgene expression in human cells (Smith et al., 2008), via CRISPR/Cas9-mediated homologous recombination (**Fig. 2A-C**). Three different CAR constructs were designed using previously-reported T or natural killer (NK) cell-specific transmembrane and intracellular domains against GBM or other solid tumors: CLTX-T-CAR (Wang et al., 2020a), interleukin 13 receptor alpha 2 (IL13Rα2)-targeted quadruple mutant IL-13 (TQM13) T-CAR (IL13-T-CAR) (Kim et al., 2020a; Nguyen et al., 2012), and CLTX-NK-CAR (Li et al., 2018) (**Fig. 2A-B**). After nucleofection, puromycin-resistant (PuroR) single cell-derived hPSC clones were isolated and subjected for PCR genotyping. Approximately 60% (3 out of 5), 53.8% (7 out of 13), and 7.7% (1 out of 13) of the clones were targeted in one allele (heterozygous), and approximately 20% (1 out of 5), 7.7% (1 out of 13) and 23.1% (3 out of 13) in both alleles (homozygous) for CLTX-T-CAR, IL-13-T-CAR, and CLTX-NK-CAR (**Fig. 2C**), respectively. While we used a relatively safe single guide RNA (sgRNA) for *AAVS1* targeting (Chen et al., 2015), off-target remains a major concern. Our sequencing results indicated no detectable insertion or deletion (indel) formation in 5 top potential off-target sites (**Table S1**), consistent with previous reports on the specificity of Cas9-mediated genome editing in hPSCs (Chen et al., 2015; Smith et al., 2014; Veres et al., 2014). The stable expression of synthetic CARs on the genetically-modified hPSCs were confirmed by RT-PCR analysis of CLTX-IgG4 and IL-13 fragments (**Fig. 2D**, **S3A**) and flow cytometry analysis of anti-IgG4 (SmP)-FITC and IL13Rα2-FITC (**Fig. 2E**) at different stages of neutrophil differentiation. Importantly, CAR-expressing hPSCs retained high expression levels of pluripotent markers SSEA-4 and OCT-4 (**Fig. S3B-C**).

**Fig. 2.**
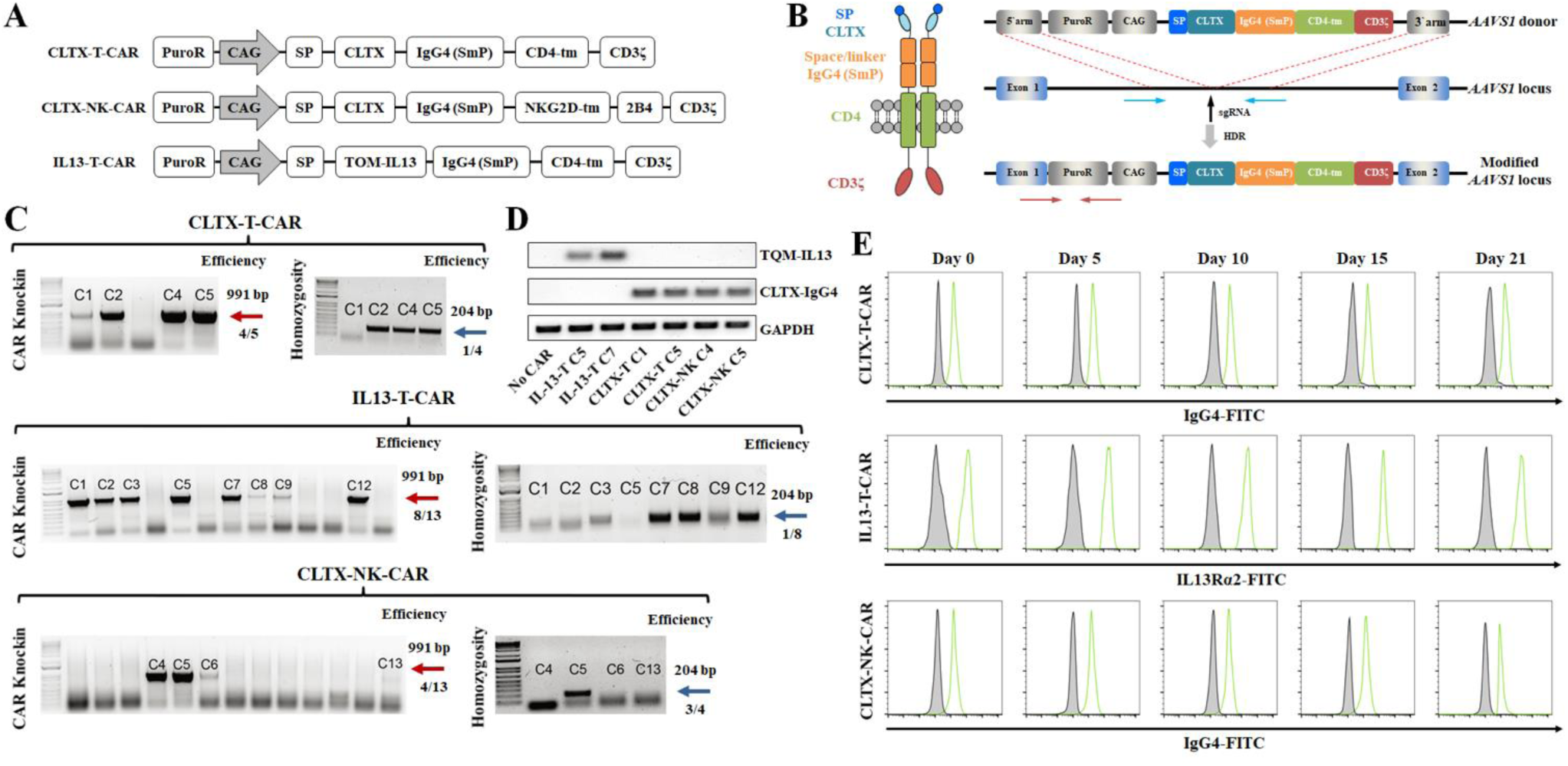
Construction of chimeric antigen receptor (CAR) knock-in H9 hPSCs using CRISPR/Cas9. (A) Schematic of CLTX-T-CAR, CLTX-NK-CAR, and IL13-T-CAR donor plasmids, composed of a signal peptide (SP), a glioblastoma-targeting extracellular domain chlorotoxin (CLTX) (Wang et al., 2020a) or quadruple mutant IL-13 (TQM13) (Kim et al., 2020a; Nguyen et al., 2012), an Fc domain IgG4 (SmP), a transmembrane domain CD4-tm, and an intracellular signal transduction domain CD3ζ. Based on a previous optimized CAR design for NK cells (Li et al., 2018), CLTX-NK-CAR also includes a 2B4 co-stimulatory domain. (B) Schematic of CLTX-T-CAR construct and targeted knock-in strategy at the AAVS1 safe harbor locus. Vertical arrow indicates the AAVS1 targeting sgRNA. Red and blue horizontal arrows indicate primers for assaying targeting efficiency and homozygosity, respectively. (C) PCR genotyping of single cell-derived hPSC clones after puromycin selection is shown, and the expected PCR product for correctly targeted AAVS1 site is 991 bp (red arrow) with an efficiency of 4 clones from a total of 5, 8 clones from a total of 13, and 4 clones from a total of 13 for CLTX-T-CAR, IL13-T-CAR, and CLTX-NK-CAR, respectively. A homozygosity assay was performed on the knock-in clones, and those without ∼240 bp PCR products were homozygous (blue arrow). (D-E) Heterozygous clones of CLTX-T-CAR C5, IL13-T-CAR C7 and CLTX-NK-CAR C5 were selected for subsequent studies. Representative RT-PCR (D) and flow cytometry (E) analysis of IL-13 and CLTX-IgG4 expression on wildtype and CAR knock-in hPSCs during neutrophil differentiation is shown.

CLTX or IL-13 CAR-expressing hPSCs presented a similar efficiency in neutrophil production as wildtype H9 hPSCs, and all the resulting neutrophils derived from different hPSCs displayed similar surface phenotypes (**Fig. 3A**). We next performed bulk RNA sequencing (RNA-seq) analysis on hPSC-derived neutrophils and compared to the transcriptome of hPSCs and healthy primary neutrophils (Perez et al., 2020). As compared to undifferentiated hPSCs, most of expression patterns of key surface markers and transcription factors were identical in different groups of hPSC-derived and primary neutrophils (**Fig. 3B-C**, **Table S2-3**). Consistent with flow cytometry analysis, RNA-seq data confirmed the expression of *ITGAM* (*CD11b*), *FUT4* (*CD15*), *FCGR3A* (*CD16*), *CEACAM8* (*CD66b*), *ITGB2* (*CD18*), and *MPO*, in hPSC-derived and primary neutrophils (Perez et al., 2020) (**Fig. 3B-C**). Notably, hPSC-derived neutrophils demonstrated a relatively low expression level of *CEACAM8* and a high expression level of *MPO*, similar to the expression levels in mature and immature primary neutrophils, respectively (**Table S2-3**). Similarly, other surface receptors, including toll-like receptors (TLRs) (Hayashi et al., 2003), adhesion molecules such as *SELL* and *ITGAX* (Rincón et al., 2018), key transcription factors such as *SPI1* (*PU.1*), *CEBPA* (*C/EBP-α*), and *CEBPE* (*C/EBP-ε*) (Sweeney et al., 2016), functional genes such as *PRTN3* and *MPO* (Lawrence et al., 2018), and genes involved in ROS production such as *NCF2* and *NCF4* (Rincón et al., 2018), are expressed at levels similar to that found in primary immature or mature neutrophils (**Fig. 3B-C**, **Table S2-3**), consistent with previous studies (Rincón et al., 2018; Sweeney et al., 2016). Despite of the similarity, significant differences between hPSC-derived and primary neutrophils are also apparent. For instance, transcription factors associated with early myeloid or granulocyte progenitors, such as *RUNX1* (Sweeney et al., 2016) and *GFI1* (Lawrence et al., 2018), retained high expression levels in hPSC-derived neutrophils, indicating an immature phenotype and/or high heterogeneity of neutrophil differentiation cultures. Chemokines and chemoattractants, including C-X-C motif chemokine receptors (CXCRs) and formyl peptide receptors (FPRs) (Rincón et al., 2018), displayed lower expression levels than intermediate and mature neutrophils, suggesting that hPSC-derived neutrophils may be less sensitive to chemoattractants than primary neutrophils. Fresh hPSC-neutrophils displayed similar expression patterns as mature primary neutrophils in terms of N1 and N2 markers (Shaul et al., 2016), (**Fig. S3D**), but unique N1 or N2 transcriptional profile was not observed in all neutrophils since they expressed a subset of N1 and N2 genes at both high and low levels (**Table S4**), highlighting the need for more authentic markers enabling the tracking of N1 or N2 human neutrophils (Perez et al., 2020). Transcriptional heterogeneity was also observed in immature, intermediate and mature primary neutrophils, indicating the dynamics and plasticity of neutrophils.

**Fig. 3.**
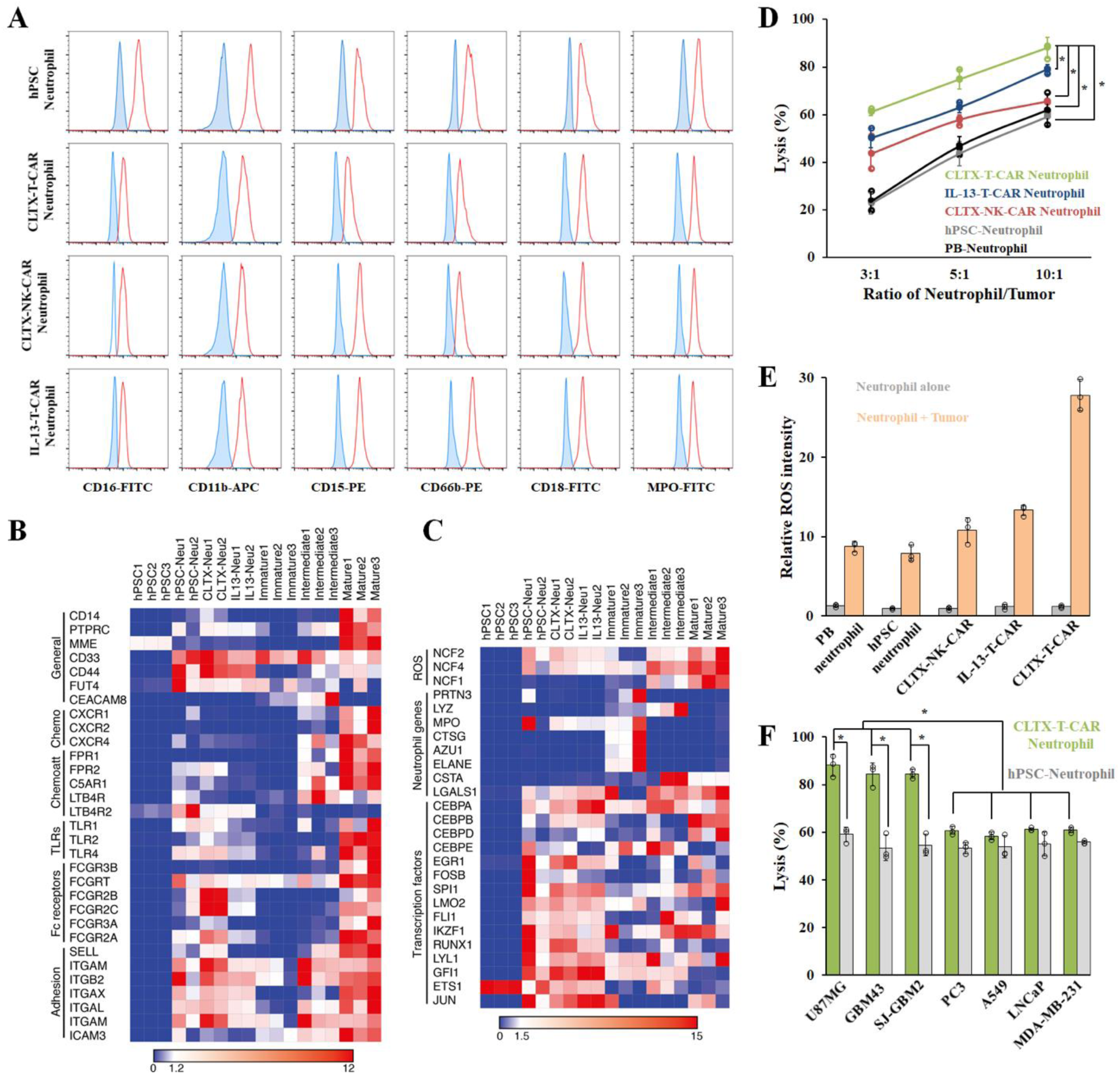
Neutrophils derived from chimeric antigen receptor (CAR) knock-in H9 hPSCs display enhanced antitumor cytotoxicity. (A) Flow cytometric analysis of neutrophils derived from different hPSCs. Plots show unstained control (blue) and specific antibody (red) histograms. (B-C) Bulk RNA sequencing analysis was performed on hPSC-derived wildtype, CLTX-T-CAR, and IL13-T-CAR neutrophils, and compared to the transcriptome of immature, intermediate and mature primary human neutrophils (Perez et al., 2020). Heatmaps show selected general surface markers, chemokines (Chemo), chemoattractants (Chemoatt), toll-like receptors (TLRs), Fc receptors, adhesion molecules (B), Reactive oxygen species (ROS) generation-related genes, other neutrophil function-related genes and transcription factors (C). (D) Cytotoxicity assays against U87MG glioblastoma were performed at different ratios of neutrophil-to-tumor target using indicated neutrophils. Data are represented as mean ± s.d. of three independent replicates, \**p*<0.05. (E) Reactive oxygen species (ROS) generation of different neutrophils co-cultured with or without U87MG cells was measured. (F) The cytotoxicity ability of CLTX-T-CAR hPSC-neutrophils against various tumor cells at a ratio of 10: 1 are shown. Glioblastoma U87MG cell line, primary adult GBM43, and pediatric SJ-GBM2 cells were employed. Data are represented as mean ± s.d. of three independent replicates. \**p*<0.05, glioblastoma versus non-glioblastoma tumor.

To determine the effects of CAR expression on the anti-tumor cytotoxicity of neutrophils, CAR-expressing hPSCs were differentiated into CAR-neutrophils (**Fig. 1A**), which were then co-cultured with glioblastoma (GBM) U87MG cells *in vitro* at different effector-to-target ratios. Consistent with enhanced cytotoxicity efficacy of CLTX (Wang et al., 2020a) and TQM13 (Kim et al., 2020a) CAR-T cells against GBM, CAR-neutrophils presented improved tumor-killing ability as compared to wildtype hPSC-derived or primary PB neutrophils (**Fig. 3D**). Among the different CAR groups, CLTX-T-CAR hPSC-neutrophils displayed superior tumor-killing activities. Neutrophils could also release cytotoxic ROS to kill target cells (Yan et al., 2014), and the kinetics of ROS production in different neutrophils coincided with their increased tumor killing abilities (**Fig. 3E**), indicating the potential involvement of ROS in neutrophil-mediated cytotoxicity against GBM cells. In addition, enhanced anti-tumor cytotoxicity was observed in the co-incubation of CLTX-T-CAR hPSC-neutrophils and GBM cells, including U87MG cell line, primary adult GBM43, and pediatric SJ-GBM2 cells (**Fig. 3F**), but not with other cancer cells, suggesting their high specificity towards GBM. Notably, CLTX-T-CAR neutrophils did not kill normal hPSCs, hPSC-derived cells, or non-tumor glial cells (**Fig. S3E-F**), consistent with a previous report that primary neutrophils do not kill healthy epithelial cells (Yan et al., 2014). Collectively, hPSC-derived CAR-neutrophils presented enhanced anti-tumor cytotoxicity against target cancer cells and produced more ROS *in vitro*.

### CLTX-T-CAR hPSC-neutrophil-mediated glioblastoma killing involves phagocytosis, ROS production and NET formation

To explore underlying mechanisms of CAR-neutrophil-mediated anti-tumor cytotoxicity, direct effector-target interactions were investigated since intimate neutrophil-tumor conjugate formation was required for cytotoxicity by neutrophils (Matlung et al., 2018). Immunological synapses between neutrophils and targeted tumor cells formed after half-an-hour co-culture and increased proportionally with incubation time (**Fig. 4A-B**). As expected, more effector-target interactions were observed between CLTX-T-CAR hPSC-neutrophils and tumor cells as compared to primary PB- and hPSC-neutrophils (**Fig. 4B**), whereas no immunological synapses formed between CAR-neutrophils and normal hPSCs, hPSC-derived cells, or non-tumor glial cells (**Fig. 4C**), highlighting their specificity against tumor cells. Live cell imaging revealed that CAR-neutrophils actively migrated toward tumor cells and uptook pre-loaded cytosolic dye Calcein-AM as early as half an hour following co-incubation (**Fig. 4D-E**). Phagocytosis of tumor cells by neutrophils was significantly reduced after treatment of 5 μM cytochalasin D (CytoD), a chemical that inhibits both phagocytosis (Esmann et al., 2010) and neutrophil extracellular trap (NET) formation (Neubert et al., 2018). Furthermore, the dynamics of ROS release agreed well with the kinetics of neutrophil phagocytosis and significantly blocked by ROS inhibitor acetylcysteine (NAC) (**Fig. 4F**). We next assessed the formation of NETs during neutrophil and tumor co-incubation in the absence or presence of NET inhibitor propofol (Meier et al., 2019). PicoGreen staining demonstrated significant decrease of NET formation in neutrophils treated with 3 μg/ml or more propofol (**Fig. 4G**). While all three inhibitors significantly blocked tumor lysis by CAR neutrophils, glioblastoma demonstrated higher viability under CytoD and NAC conditions (**Fig. 4H**). Consistent with previous reports (Matlung et al., 2018; Yan et al., 2014), our results demonstrated that tumor-killing of hPSC-derived neutrophils involves phagocytosis, ROS release and NET formation.

**Fig. 4.**
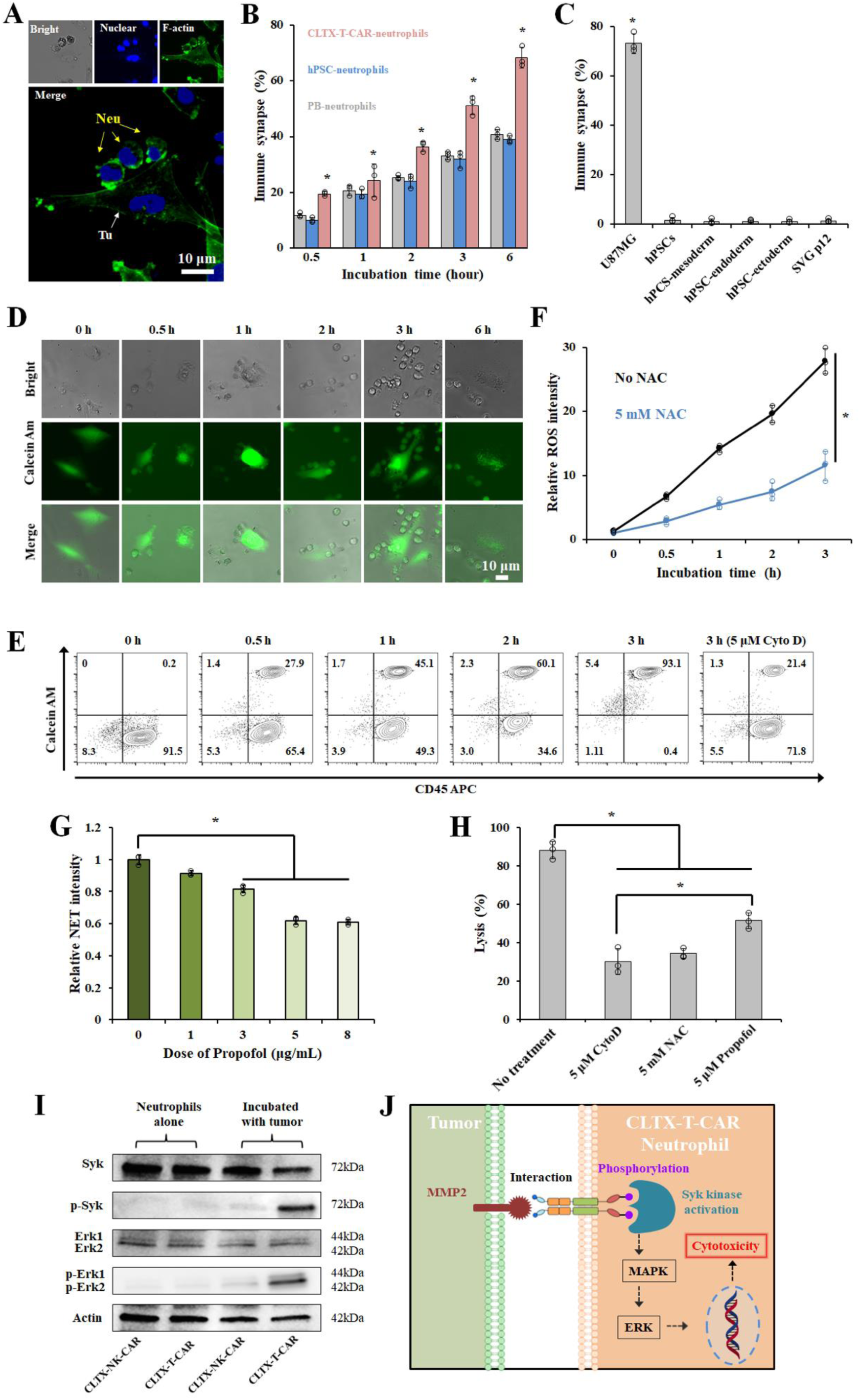
CLTX-T-CAR neutrophil-mediated tumor lysis involves phagocytosis, ROS production and NET formation. (A-C) Representative images of immunological synapses indicated by polarized F-actin accumulation at the interface between CAR-neutrophils and tumor cell were shown in (A) and the numbers of immunological synapses formed between indicated neutrophils and tumor cells were quantified in (B). Neu: neutrophils; Tu: tumor cells. Scale bars, 10 μm. (C) The numbers of immunological synapses formed between CLTX-T-CAR hPSC-neutrophils and indicated cells were quantified. Data are represented as mean ± s.d. of three independent replicates. (D-E) Phagocytosis of glioblastoma cells by neutrophils led to a reduction in cancer cell cytoplasmic labeling in a time-dependent manner. Phagocytosis was significantly blocked when neutrophils were treated with 5 μM cytochalasin D (CytoD). Representative brightfield (bright) and fluorescent images of the phagocytotic disruption of Calcein-AM-labeled tumor cells (D), and flow cytometric analysis of CLTX-T-CAR hPSC-neutrophils during tumor phagocytosis (E) were shown. Scale bars, 10 μm. (F) Reactive oxygen species (ROS) generation during the phagocytotic disruption of tumor cells by CLTX-T-CAR hPSC-derived neutrophils with or without 5 mM acetylcysteine (NAC) was quantified. (G) Formation of neutrophil extracellular traps (NETs) in CLTX-T-CAR neutrophils treated with indicated doses of propofol was quantified using PicoGreen. (H) Tumor lysis of CLTX-T-CAR neutrophils treated with or without 5 μM CytoD, 5 mM NAC and 5 μM propofol was quantified. (I) Total and phospho-protein analysis of the Syk-Erk signaling pathway in cell lysates of CLTX-T-CAR and CLTX-NK-CAR hPSC-neutrophils via western blot was performed with or without co-incubation of tumor cell. (J) Schematic of activated cellular Syk-Erk signaling pathway in CAR-neutrophils after binding to MMP2.

### CLTX-T-CAR hPSC-neutrophils specifically bind to glioblastoma via MMP2

To further explore molecular mechanisms underlying CLTX-T-CAR-enhanced cytotoxicity against tumor cells, we determined the biological function of potential CLTX ligands, including chloride channels (CLCN3), phospholipid protein annexin A2 (ANXA2), and matrix metalloproteinase 2 (MMP2) as previously reported (Wang et al., 2020a). We used an inducible Cas13d-mediated gene knockdown platform (**Fig. S4A**) to suppress the candidate gene expression. After puromycin selection, approximately 78% of the transfected glioblastoma cells expressed Cas13d as indicated by eGFP expression (**Fig. S4B-C**) in the presence of doxycycline (DOX). RT-PCR analysis confirmed the successful knockdown of *CLCN3* (**Fig. S4D**), *ANXA2* (**Fig. S4E**), and *MMP2* (**Fig. S4F**) in U87MG GBM cells. Similar to the scramble control in shRNA system (Lian et al., 2013), we used non-targeting Cas13d single guide RNA (sgRNA) as a negative control (*e.g.*, *CLCN3* and *AXAN2* sgRNAs were used as negative controls for *MMP2* sgRNAs), and no obvious cross-gene knockdown effects were observed (**Fig. S4G**). Notably, knockdown of *MMP2*, but not *CLCN3* or *ANXA2*, significantly reduced CLTX-T-CAR neutrophil-mediated tumor cell killing (**Fig. S4F**), consistent with a previous report on CLTX CAR-T cells (Wang et al., 2020a). To further determine the relationship between CAR-neutrophil activity and *MMP2*, we assessed gene expression levels of *ANXA2*, *CLCN3* and *MMP2* in different tumor cells. As expected, U87MG, GBM43 and SJ-GBM cells displayed highest expression levels of *MMP2* (**Fig. S4H**). Linear regression analysis demonstrated tumor lysis of CLTX-T-CAR neutrophils is most likely dependent on the *MMP2* expression (**Fig. S4I-K**). As compared to GBM cells, *MMP2* is expressed at a low to negligible level in normal SVG p12 glial cells and hPSC-derived somatic cells (**Fig. S4L**), consistent with their minimal apoptosis in CAR-neutrophil-mediated lysis (**Fig. 4C**). Consistent with a previous report on CLTX CAR-T cells (Wang et al., 2020a), these findings further demonstrate that membrane protein MMP2 is required for CLTX-T-CAR recognition and activation of CAR-neutrophils to kill tumor cells. This also suggests the safety of CLTX-T-CAR neutrophils in future clinical application, given the low or negligible MMP2 expression on human normal tissues as compared to GBM (Itoh, 2015; Lyons et al., 2002).

We next investigated downstream intracellular signaling in activated neutrophils after binding to MMP2-expressing tumor cells. Primary neutrophils display antibody-dependent cellular cytotoxicity (ADCC) toward tumor cells via trogoptosis, which is mediated by Fcγ receptor and its downstream signaling pathways, including tyrosine kinase Syk (Matlung et al., 2018). CLTX-T-CAR hPSC-neutrophils displayed stronger phosphorylated activation of Syk (p-Syk) upon GBM stimulation, compared to their counterparts with CLTX-NK-CAR (**Fig. 4I**). Notably, significantly increased ratio of extracellular signal-regulated kinase (Erk) 1/2 (p-Erk1/2), a key signaling mediator involved in lymphoid-mediated cytotoxicity (Li et al., 2018), was also observed in our GBM-stimulated CLTX-T-CAR hPSC-neutrophils. This indicates potential activation of Syk-vav1-Erk pathway in activated neutrophils (**Fig. 4J**), reminiscent of intracellular signaling transduction in CAR hPSC-NK cells (Li et al., 2018).

### CLTX-T-CAR hPSC-neutrophils display high transmigration and anti-tumor cytotoxicity activities in biomimetic tumor models *in vitro*

To further evaluate the activities of CAR-neutrophils, we implemented a transwell-based blood brain barrier (BBB) model (**Fig. 5A**) using human cerebral microvascular endothelial cells (**Fig. S4M**). While CAR-expressing and wildtype hPSC-neutrophils displayed similar transmigration activity across the BBB in response to N-Formylmethionine-leucyl-phenylalanine (fMLP) (**Fig. 5B**), CAR-neutrophils demonstrated higher tumor-killing ability than wildtype neutrophils (**Fig. 5C**). Furthermore, CLTX-T-CAR hPSC-neutrophils retained high transmigration ability during their second trafficking across the BBB in response to the inflammatory tumor cells (**Fig. 5D-5E**), recapitulating many aspects of *in vivo* inflammation and cancer. A three-dimensional (3D) GBM model was also implemented to construct an *in vivo* tumor niche-like microenvironment that contains a dense extracellular matrix network and heterogeneous tumor cell subtypes (**Fig. 5F**). As compared to the wildtype control, CLTX-T-CAR hPSC-neutrophils exhibited higher tumor-infiltrating in the 3D tumor model (**Fig. 5G**) and tumor-killing activities (**Fig. 5H**) under both hypoxic (3% O_2_) and normoxic (21% O_2_) conditions. Pre-treating neutrophils with soluble MMP2 and cytochalasin D (CytoD) significantly reduced tumor-infiltrating activity and antitumor cytotoxicity of CAR-neutrophils (**Fig. 5G, S4N-O**), consistent with our observation in monolayer cell cultures. To further explore the molecular mechanisms underlying anti- and pro-tumor activities of wildtype and CAR-neutrophils under hypoxia (Chan et al., 2021; Huang et al., 2018), a common feature of solid tumor microenvironment that contributes to immunosuppression (Rankin et al., 2016) and shapes the anti-tumor (N1) or pro-tumor (N2) phenotype of infiltrated neutrophils (Fridlender et al., 2009; Shaul et al., 2016), we performed N1 or N2 phenotype analysis on the isolated neutrophils. Compared with normoxia, hypoxia significantly decreased expression of N1-specific markers, including *ICAM-1*, *iNOS*, *TNFα* and *CCL3*, and increased N2-specific markers, including *CCL2*, *VEGF*, *CCL5*, and *Arginase* (Shaul et al., 2016), in wildtype hPSC-neutrophils (**Fig. 5I-J**). On the contrary, CLTX-T-CAR hPSC-neutrophils retained high expression levels of N1 markers under hypoxia. We next used ELISA to detect human cytokine production release in the media after neutrophil-tumor co-culture, including tumor necrosis factor-α (TNFα) and IL-6 (**Fig. 5K**). Both wildtype and CAR-neutrophils produced TNFα and IL-6 after tumor stimulation, and CAR-neutrophils maintained highest levels of both cytokines under hypoxia and normoxia. Notably, hypoxia significantly reduced cytokine release in wildtype neutrophils. Taken together, CLTX-T-CAR hPSC-neutrophils sustained an anti-tumor phenotype, and retained high transmigration ability and anti-tumor cytotoxicity under tumor-niche mimicking hypoxic conditions, highlighting their potential application in targeted immunotherapy.

**Fig. 5.**
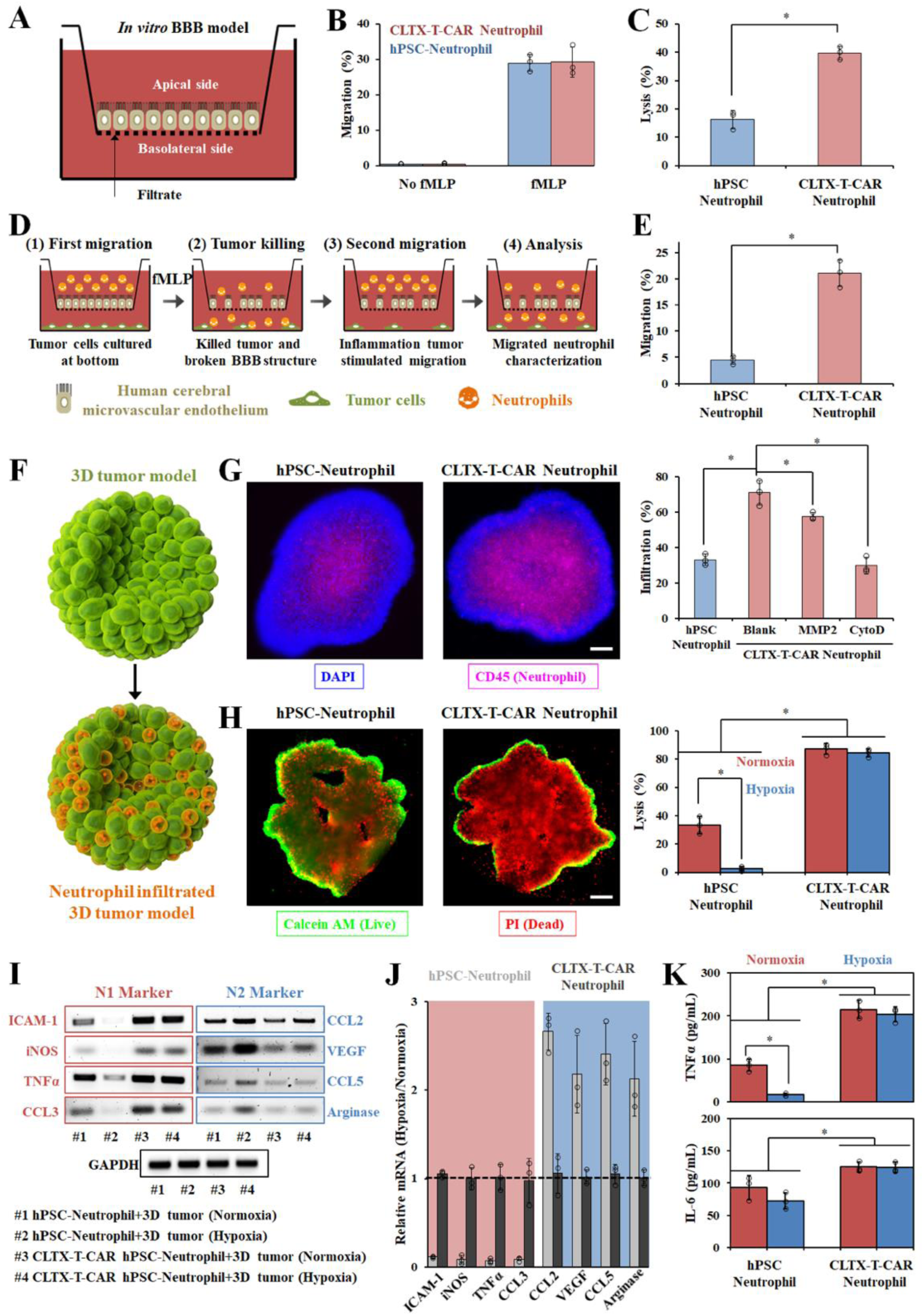
Functional evaluation of CLTX-T-CAR hPSC-neutrophils using glioblastoma (GBM) microenvironment mimicking models *in vitro*. (A) Schematic of *in vitro* blood-brain-barrier (BBB) model. Transwell migration analysis of wildtype and CLTX-T-CAR hPSC-neutrophils with or without 100 nM of fMLP treatment (B), and their anti-GBM cytotoxicity (C) were assessed in the BBB model. Schematic (D) and quantification (E) of second migration of different hPSC-neutrophils through BBB were shown. (F) Schematic of neutrophil-infiltrated three-dimensional (3D) tumor model *in vitro*. (G) Representative fluorescent images and quantification of infiltrated wildtype and CLTX-T-CAR hPSC-neutrophils in the 3D tumor models were shown. DAPI was used to stain the cell nuclear and CD45 was used to stain neutrophils. Data are represented as mean ± s.d. of three independent replicates, \**p*<0.05. (H-K) Phenotype analysis of tumor-infiltrated neutrophils was performed under normoxia or hypoxia. (H) Live/dead staining of the 3D tumor model was performed after 24 hr of neutrophil infiltration and the corresponding tumor-killing efficiency was quantified using cytotoxicity kit. Data are represented as mean ± s.d. of three independent replicates, \**p*<0.05. Scale bars, 200 μm. (I) Wildtype and CLTX-T-CAR hPSC-neutrophils were isolated from the 3D tumor models and subjected for RT-PCR analysis of anti-tumor N1 and pro-tumor N2 markers. (J) Quantification of the relative mRNA intensity for anti-tumor N1 and pro-tumor N2 markers was shown. Data are mean ± s.d. in (I). (K) Cytokine production release in the media after neutrophil-tumor co-culture, including tumor necrosis factor-α (TNFα) and IL-6, was measured by ELISA.

### CLTX-T-CAR hPSC-neutrophils display enhanced activity against glioblastoma *in vivo*

To determine the function of CLTX-T-CAR hPSC-neutrophils *in vivo*, we implemented an *in situ* xenograft model via intracranial injection of 5×10^5^ luciferase-expressing GBM cells into the brain of immunodeficient mice (Wang et al., 2020b). Neutrophils were administrated intratumorally or intravenously to investigate their *in vivo* tumor-killing activities, in comparison to hPSC-derived CAR-natural killer (NK) cells (**Fig. S5A-E**). Notably, CLTX-T-CAR neutrophils are more effective in killing GBM cells than CLTX-NK-CAR NK cells *in vitro*, and the combinatory effect between CAR-neutrophils and CAR-NK cells was not observed. In the intratumoral injection experiment, tumor-bearing mice were intratumorally administrated with a single dose of PBS, 5×10^6^ CLTX-T-CAR hPSC-neutrophils, CLTX-NK-CAR hPSC-NK cells, or corresponding wildtype controls 3 hour following tumor cell inoculation (**Fig. S5F**). Bioluminescent imaging (BLI) was performed to monitor tumor growth weekly after initial imaging on day 3 (**Fig. S5G**). As compared to PBS-treated mice, treatment with hPSC-derived neutrophils or NK cells significantly reduced tumor burden (**Fig. S5G-H**). As expected, hPSC-derived CLTX-NK-CAR NK cells and CLTX-T-CAR neutrophils displayed higher anti-tumor cytotoxicity than their wildtype controls in the mice that maintained a stable body weight (**Fig. S5I**). Notably, one of the PBS-treated tumor-bearing mice died at day 30 due to the overgrowth of tumor in the recipient brain (**Fig. S5G**, **S5J**).

We next investigated *in vivo* activities of CAR-neutrophils via weekly intravenous administration of 5×10^6^ neutrophils into tumor-bearing mice (**Fig. 6A**). To track *in vivo* biodistribution and trafficking of CAR-neutrophils, we labeled neutrophils with Cy5 before systemic injection and performed fluorescence imaging 1, 5 and 24 hours after systemic administration (**Fig. S6A**). Neutrophils trafficked to the whole mouse body in an hour and retained a similar biodistribution in 5 hours after neutrophil injection (**Fig. S6B**). Compared to wildtype neutrophils, CAR-neutrophils effectively crossed BBB and trafficked to GBM xenograft in the mouse brain after 24 hours (**Fig. S6C-D**). No significant changes of body weight were observed across the experimental mouse groups during the intravenous administration study (**Fig. S6E**). Consistent with the intratumoral administration study, CAR-neutrophils displayed higher anti-tumor cytotoxicity than PBS and wildtype controls in mice according to BLI analysis (**Fig. 6B-C**) as well as brightfield and H&E staining images of GBM xenografts (**Fig. S6F**). Notably, tumor-bearing mice treated with CLTX-T-CAR hPSC-neutrophils demonstrated a significantly reduced tumor burden as compared to those treated with CAR-NK cells, suggesting the superior ability of neutrophils in crossing BBB and penetrating GBM xenograft in mice. In contrast to CAR-neutrophils, weekly administration of wildtype hPSC-neutrophils or peripheral blood (PB) neutrophils significantly promoted the growth of tumor in the brain with or without CAR-NK cells, and resulted in the death of tumor-bearing mice as early as day 21 (**Fig. 6D**). Despite the rarity of extraneural metastasis of GBM (∼200 reported cases (Rosen et al., 2018)) and unknown pathogenesis (Seo et al., 2012), systemic metastasis occurred in the tumor-bearing mice treated with wildtype hPSC-derived or PB-neutrophils as determined by BLI images of various *ex vivo* organs and/or tissues (**Fig. 6E-F**), suggesting a potential role of neutrophils in the extracranial metastasis of human patient GBM (Liang et al., 2014; Wang et al., 2020c).

**Fig. 6.**
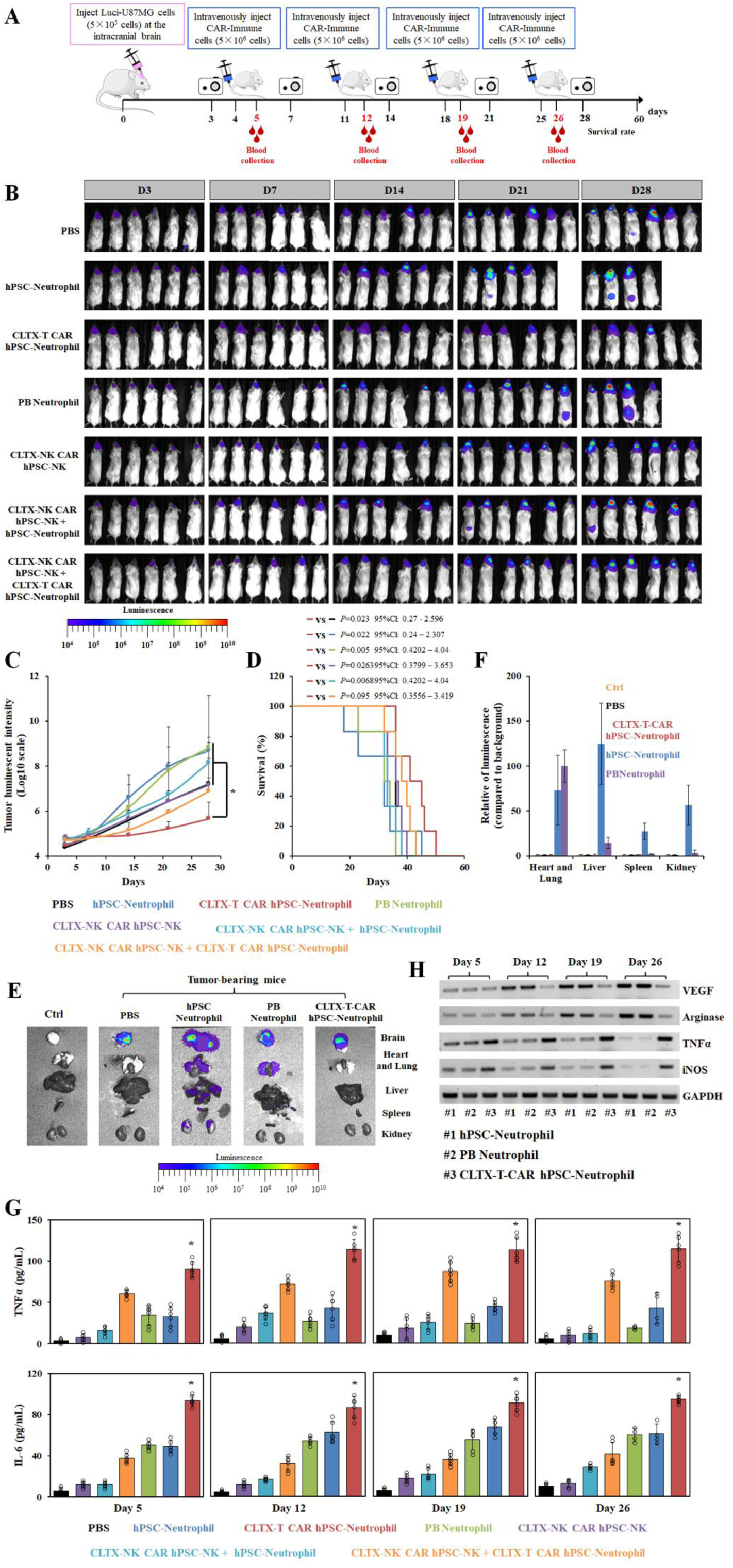
In vivo anti-tumor activities of hPSC-derived CLTX-T-CAR neutrophils and CLTX-NK-CAR NK cells were assessed via intravenous injection. (A) Schematic of intravenous injection of CAR-neutrophils and/or CAR-NK cells for *in vivo* anti-tumor cytotoxicity study. 5×10^5^ luciferase (Luci)-expressing U87MG cells were stereotactically implanted into the right forebrain of NSG mice. After 4 days, mice were intravenously treated with PBS, 5×10^6^ wildtype neutrophil, CAR-neutrophils, wildtype NK cells, and/or CAR-NK cells weekly for about a month. Time-dependent tumor burden was determined (B) and quantified (C) by bioluminescent imaging (BLI) at the indicated days. Data are mean ± s.d. for the mice in (B) (n=6). (D) Kaplan-Meier curve demonstrating survival of the experimental groups was shown. Organs harvested from dead mice were subjected for bioluminescent imaging (E) and quantified in (F). Data are mean ± s.d. for the mice in (E) (n=6). (G) Levels of human tumor necrosis factor-α (TNFα) and IL-6 in mouse peripheral blood were measured by ELISA. (H) Wildtype and CAR-neutrophils were isolated from mouse blood 24 hour after systemic injection, and subjected for RT-PCR analysis of anti-tumor N1 and pro-tumor N2 markers.

We next measured human cytokine production release in plasma of different experimental mouse groups, including TNFα and IL-6. All non-PBS experimental groups produced detectable TNFα and IL-6 in plasma from day 5 to day 26, and CAR-neutrophils maintained highest levels of both cytokines (**Fig. 6G**). To further explore underlying mechanism of neutrophil-mediated metastasis, we harvested human neutrophils from mouse blood and subjected for anti-tumor (N1) or pro-tumor (N2) phenotype analysis (Fridlender et al., 2009; Shaul et al., 2016). Tumor xenografts significantly decreased expression of N1-specific markers, including *iNOS* and *TNFα*, and increased N2-specific markers, including *VEGF* and *Arginase* (Shaul et al., 2016), in wildtype hPSC- or PB-neutrophils (**Fig. 6H**, **S6G**). On the contrary, CLTX-T-CAR hPSC-neutrophils retained high expression levels of N1 markers, which is consistent with their strong anti-tumor cytotoxicity and cytokine release in tumor-bearing mice. Collectively, our findings clearly demonstrated that hPSC-derived CAR-neutrophils can sustain an anti-tumor phenotype and efficiently kill tumor cells under various tumor niche-like conditions tested in this study, highlighting their potential application in targeted immunotherapy.

## DISCUSSION

While immunotherapy has been developed to cure hematologic malignancies, *de novo* and acquired resistance to targeted cancer therapy is commonly observed in solid tumors due to the complex and dynamic tumor microenvironment (TME), in which neutrophils are key players (Devlin et al., 2020; Kalafati et al., 2020; Ponzetta et al., 2019). Increased understanding of neutrophil contributions to the TME has increased interest in reprograming and/or depleting pro-tumor neutrophils as an alternative approach to treat cancer (Kalafati et al., 2020). In contrast to previous neutrophil depletion approaches, we demonstrate here the feasibility of using synthetic CARs to program and maintain neutrophils as anti-tumor effector cells both *in vitro* and *in vivo*, representing an advanced neutrophil-based immunotherapy that may complement current standard cancer treatments and boost their efficacy.

Due to the short-life of primary neutrophils and their resistance to genome editing, engineering hPSCs with synthetic CARs should be an ideal approach to produce off-the-shelf CAR-neutrophils. To achieve this goal, we first developed a chemically-defined, feeder-free platform for robust production of neutrophils from hPSCs, using stage-specific employment of signaling pathway modulators. Based on previous studies on CAR-T (Wang et al., 2020a) and CAR-NK cells (Li et al., 2018), we designed and evaluated three different CAR constructs (**Fig. 2A**) with NK or T cell-specific transmembrane and intracellular activation domains in enhancing neutrophil-mediated tumor killing activities. CLTX-T-CAR, that contains a GBM-binding peptide chlorotoxin and T cell-specific signaling domains (Wang et al., 2020a), markedly improved antigen-specific tumor cytotoxicity of hPSC-neutrophils to a level comparable to CAR-T cells with a similar CLTX-T-CAR (Wang et al., 2020a) and superior than that of CAR-macrophages with an anti-CD19 CAR *in vitro*(Klichinsky et al., 2020), though no quantitative comparison data is available in mice due to the different *in vivo* tumor models. Notably, systematically administered CAR-neutrophils presented superior anti-GBM activities in mice as compared to hPSC-derived CAR-NK cells, possibly due to a better ability of neutrophils to cross BBB and penetrate GBM xenografts in mice. In future studies, it will be interesting to investigate whether neutrophil-specific transmembrane and activation domains can be harnessed to establish neutrophil-specific CAR constructs (Roberts et al., 1998). Using an inducible gene knockdown system, we identified membrane protein MMP2 on GBM cells as the target of CLTX-binding and recognition that triggers CAR activation in neutrophils. Molecular mechanism investigation revealed that CLTX-T-CAR triggers known downstream intracellular signaling pathways that mediate phagocytosis activity against tumor cells. Gene expression profiles in neutrophils indicated a sustained anti-tumor N1 phenotype under various tumor niche-like conditions.

In summary, the CAR-neutrophil engineering platform described in this study can serve as a scalable strategy to make off-the-shelf neutrophils as potential standardized cellular products for clinical applications in cancer and neutropenia treatment. Given the relative ease of genome editing in hPSCs, other genetic modifications, such as multiple CAR expression and/or inhibitory receptor deletions, can also be performed to achieve optimal therapeutic effects in CAR-neutrophils. Due to their native ability to cross the BBB and penetrate the brain parenchyma, neutrophil-mediated delivery of imaging and chemotherapeutic drugs into brain has improved the diagnosis and treatment of GBM (Wu et al., 2018; Xue et al., 2017), and such a combination may further enhance antitumor activities of hPSC-derived CAR-neutrophils. Importantly, stable CAR-expressing hPSC lines can be also used to produce off-the-shelf CAR-T and -NK cells such as those that are currently used in clinical trials (Li et al., 2018).

### Limitations of study

The U87MG cell line used in this study provides a proof-of-concept of anti-tumor activity of hPSC-derived CAR-neutrophils, but these findings need validation in models that represent the heterogeneity of cells and tumors found in human patients (Wang et al., 2020a). In addition, the effects of neutrophils on lymphoid infiltration and immuno-landscape in GBM tumor have not been examined in this study due to our limited access to suitable humanized animal models of GBM. The combinatorial administration of CAR-neutrophils, T and NK cells into NSG mice may provide more insight into the dynamics of tumor niche in GBM after neutrophil injection. The implementation of preclinical large animal studies focusing on safety and survival will be important to better evaluate the therapeutic effects of CAR-neutrophils following multidisciplinary clinical treatment with maximal surgical resection and radiotherapy/chemotherapy.

## ACKNOWLEDGEMENTS

We thank members of the Deng and Bao laboratories for technical assistance and critical reading of the manuscript, and Dr. Sean P. Palecek for valuable discussion and advice. We’re also gratefully acknowledge the Purdue Flow Cytometry and Cell Separation Facility, Purdue Genomics Core Facility and the Biological Evaluation Core at Purdue University Center for Cancer Research (PCCR). We also thank Dr. Sandro Matosevic for providing us the primary glioblastoma cells, and Dr. Edouard G. Stanley and Andrew G. Elefanty for the H9 SOX17-mCherry/RUNX1c-GFP dual reporter line. This study was supported by startup funding from the Davidson School of Chemical Engineering and the College of Engineering at Purdue (X.B.), PCCR Robbers New Investigators (X.B.), Showalter Research Trust (Young Investigator Award to X.B.), NIH NIGMS (grant no. R35GM119787 to Q.D.), NIH NHLBI (grant no. R35HL139599 to H.E.B.), and NIH NIDDK (grant no. U54DK106846). The authors also gratefully acknowledge support from the Purdue University Center for Cancer Research, P30CA023168, Purdue Institute for Integrative Neuroscience (PIIN) and Bindley Biosciences Center, and the Walther Cancer Foundation.

## AUTHOR CONTRIBUTIONS

Y.C. and X.B. conceived and designed most of the experiments. Y.C., G.J., S.N.H., T.W., R.S., X.L., and Q.D. designed and conducted *in vitro* functional characterization of neutrophils. Y.C., S.T., B.D.E., and X.B. designed and performed the mouse experiments. Y.C., X.W., and H.E.B. designed and performed the hypoxia-related experiments. Y.C. and X.B. wrote the manuscript with support from all authors.

## STAR★METHODS

### KEY RESOURCE TABLE

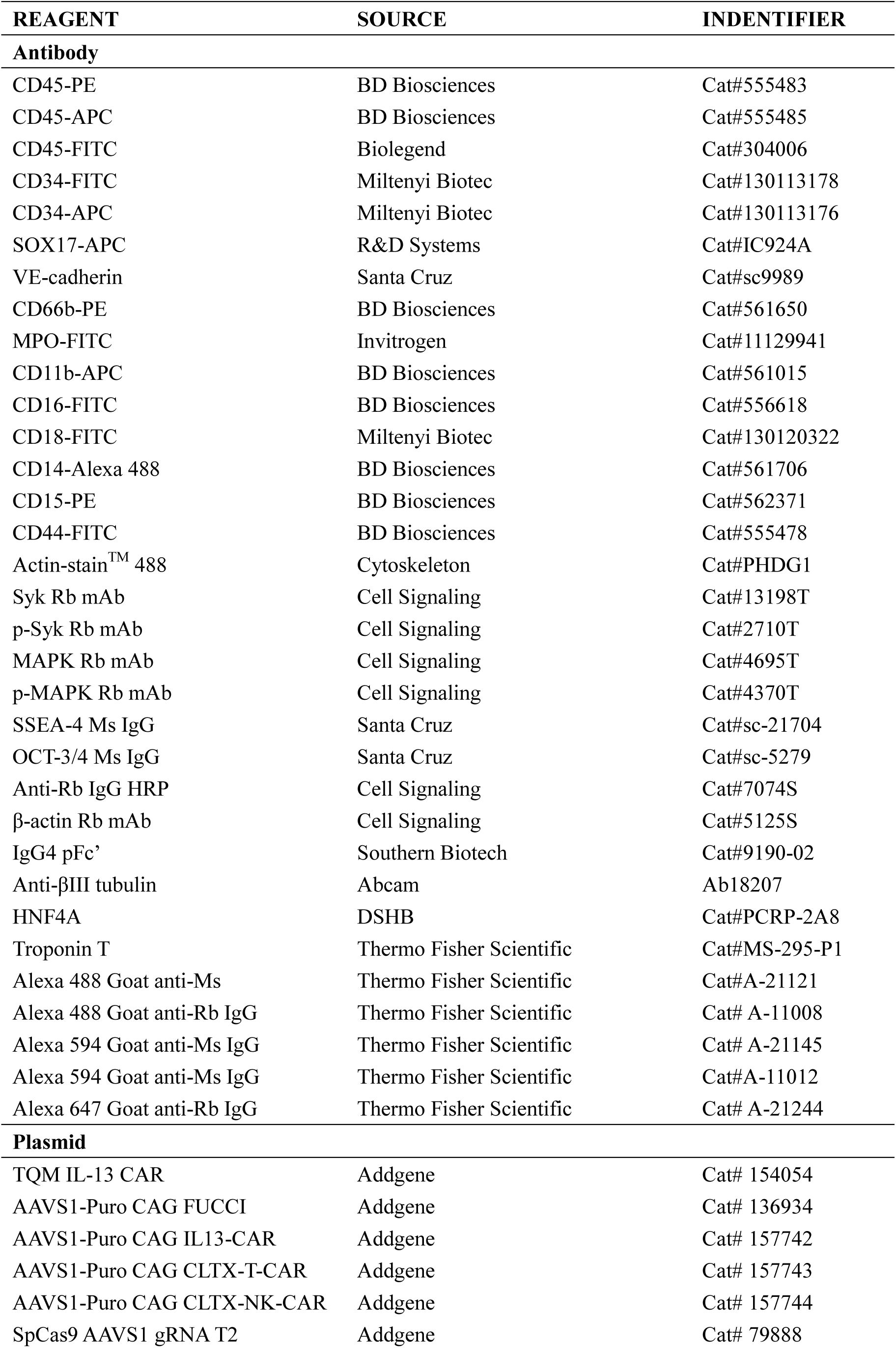
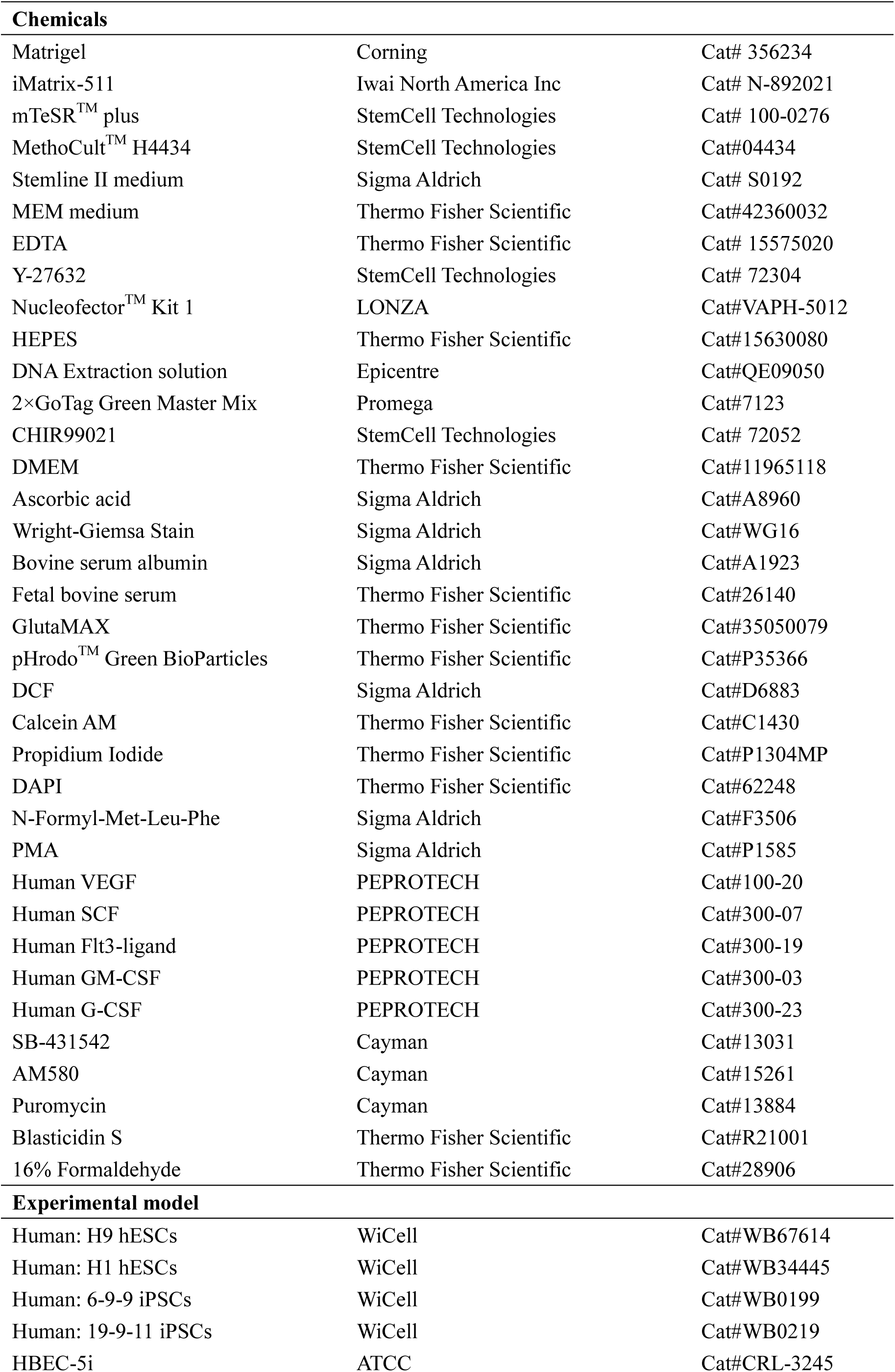
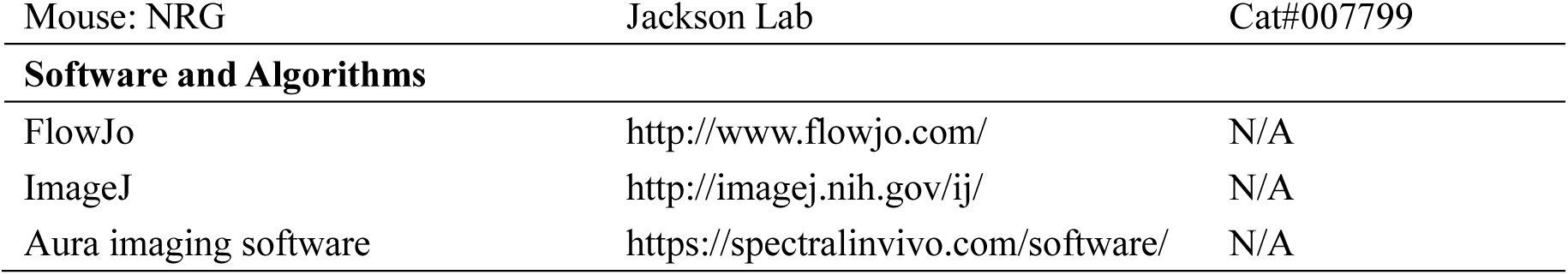

### RESOURCE AVAILABILITY

#### Lead Contact

Further information and requests for resources and reagents should be directed to, and will be fulfilled by, the Lead Contact, Xiaoping Bao.

#### Materials Availability

Human pluripotent stem cell lines H9, H1, 6-9-9 and 19-9-11 were obtained from WiCell, and the CAR hPSC lines generated in this study are available with required material transfer agreement.

#### Data and code availability

RNA-sequencing datasets generated during this study are available at NCBI GEO with accession number: GSE188393.

### EXPERIMENTAL MODEL AND SUBJECT DETAILS

#### GBM xenograft mouse models

All the mouse experiments were approved by the Purdue Animal Care and Use Committee (PACUC). The immunodeficient NOD.Cg-*RAG^1tm1Mom^IL2rg^tm1Wjl^*/SzJ (NRG) mice were bred and maintained by the Biological Evaluation Core at the Purdue University Center for Cancer Research. *In situ* xenograft murine models were constructed via intracranial injection of 5×10^5^ luciferase-expressing GBM cells into the brain of immunodeficient mice. For intratumoral administration, 5×10^6^ neutrophils or 5×10^6^ NK cells were injected after 3 days of tumor cells injection to evaluate their in vivo anti-tumor activities. For intravenous administration, 5×10^6^ neutrophils, 5×10^6^ NK cells + Neutrophils (ratio of 1:1), or 5×10^6^ NK cells were intravenously injected at day 4, day 11, day 18, and day 25. And blood was collected from these groups at day 5, day 12, day 19, and day 26. Tumor burden was monitored by bioluminescence imaging (BLI) system (Spectral Ami Optical Imaging System) and weight body of experimental mice was measured about once per week. The collected blood was stained with CD45 and analyzed in the Accuri C6 plus flow cytometer (Beckton Dickinson) after washing with PBS-/- solution containing 0.5% BSA. The collected blood was further analyzed by enzyme-linked immunosorbend assay (ELISA) for both TNFα and IL-6 (Invitrogen). At the end of treatment, tumors were collected for H&E staining. For *in vivo* biodistribution analysis, fluorescence images were captured by Spectral Ami Optical Imaging System after Cy5 (Lumiprobe) labeled neutrophils intravenously injected 1, 5, and 24 hours.

### METHOD DETAILS

#### Donor plasmid construction

The donor plasmids targeting *AAVS1* locus were constructed as previously described (Chang, et al., 2020). Briefly, to generate the CAG-IL13 T-CAR plasmid, the TQM-IL13 CAR fragment (Kim et al., 2020b) was amplified from Addgene plasmid #154054 and then cloned into the AAVS1-Puro CAG-FUCCI donor plasmid (Addgene; #136934), replacing the FUCCI. For CAG-CLTX T-CAR plasmid, the chlorotoxin sequence containing a signal peptide was directly synthesized (GeneWiz) and used to replace the IL-13 sequence in CAG-IL13 T-CAR. For CAG-CLTX NK-CAR plasmid, the conjugated NKG2D, 2B4 and CD3-ζ sequence was directly synthesized and used to replace the CD4tm and CD3-ζ sequence in CAG-CLTX T-CAR. All CAR constructs were sequenced and submitted to Addgene (#157742, #157743 and #157744).

#### Maintenance and differentiation of hPSCs.

H9, 6-9-9 and 19-9-11 hPSC lines were obtained from WiCell and maintained on Matrigel- or iMatrix 511-coated plates in mTeSR plus medium. For neutrophil differentiation, hPSCs were dissociated with 0.5 mM EDTA and seeded onto iMatrix 511-coated 24-well plate in mTeSR plus medium with 5 μM Y27632 for 24 hours (day -1). At day 0, cells were treated with 6 μM CHIR99021 (CHIR) in DMEM medium supplemented with 100 μg/mL ascorbic acid (DMEM/Vc), followed by a medium change with LaSR basal medium from day 1 to day 4. 50 ng/mL VEGF was added to the medium from day 2 to day 4 for female hPSC lines. At day 4, medium was replaced by Stemline II medium (Sigma) supplemented with 10 μM SB431542, 25 ng/mL SCF and FLT3L. On day 6, SB431542-containing medium was aspirated and cells were maintained in Stemline II medium with 50 ng/mL SCF and FLT3L. At day 9 and day 12, the top half medium was aspirated and changed with 0.5 ml fresh Stemline II medium containing 50 ng/mL SCF, 50 ng/mL FLT3L and 25 ng/mL GM-CSF. Day 15 floating cells were gently harvested and filtered for terminal neutrophil differentiation in Stemline II medium supplemented with 1X GlutaMAX, 150 ng/mL G-CSF, and 2.5 μM retinoic acid agonist AM580. Half medium change was performed every 3 days, and mature neutrophils could be harvested for analysis starting from day 21. For NK cell differentiation, day 15 floating hematopoietic stem and progenitor cells were gently harvested, filtered with a cell strainer, and cultured on OP9 DLL4 (kindly provided by Dr. Igor Slukvin) monolayer (2×10^4^ cells/mL) in NK cell differentiation medium: α-MEM medium supplemented with 20% FBS, 5 ng/mL IL-7, 5 ng/mL FTL3L, 25 ng/mL SCF, 5 ng/mL IL-15, and 35 nM UM171. NK cell differentiation medium was changed every 3 days, and the floating cells were transferred onto fresh OP9 DLL4 monolayer every 6 days.

#### Nucleofection and genotyping of hPSCs

To increase cell viability, 10 μM Y27632 was used to treat hPSCs 3–4 hr before nucleofection or overnight. Cells were then singularized by Accutase for 8-10 min, and 1-2.5 ×10^6^ hPSCs were nucleofected with 6 μg SpCas9 *AAVS1* gRNA T2 (Addgene; #79888) and 6 μg CAR donor plasmids in 100 μl human stem cell nucleofection solution (Lonza; #VAPH-5012) using program B-016 in a Nucleofector 2b. The nucleofected cells were seeded into one well of a Matrigel-coated 6-well plate in 3 ml pre-warmed mTeSR plus or mTeSR1 with 10 μM Y27632. 24 hr later, the medium was changed with fresh mTeSR plus or mTeSR1 containing 5 μM Y27632, followed by a daily medium change. When cells were more than 80% confluent, drug selection was performed with 1 μg/ml puromycin (Puro) for 24 hr. Once cells recovered, 1 μg/ml Puro was applied continuously for about 1 week. Individual clones were then picked using a microscope inside a tissue culture hood and expanded for 2–5 days in each well of a 96-well plate pre-coated with Matrigel, followed by a PCR genotyping. The genomic DNA of single clone-derived hPSCs was extracted by scraping cells into 40 μl QuickExtractTM DNA Extraction Solution (Epicentre; #QE09050). 2×GoTaq Green Master Mix (Promega; #7123) was used to perform the genomic DNA PCR. For positive genotyping, the following primer pair with an annealing temperature Tm of 65°C was used: CTGTTTCCCCTTCCCAGGCAGGTCC and TCGTCGCGGGTGGCGAGGCGCACCG. For homozygous screening, we used the following set of primer sequences: CGGTTAATGTGGCTCTGGTT and GAGAGAGATGGCTCCAGGAA with an annealing temperature Tm of 60°C.

#### Hematopoietic colony forming and Wright-Giemsa staining

Collected cells were grown in 1.5 mL of cytokine containing MethoCult H4434 medium (StemCell Technologies, Vancouver) at 37 °C. The hematopoietic colonies were scored for colony forming units (CFUs) according to cellular morphology. To assess the morphology of cells, neutrophils were fixed on glass slides and stained with Wright-Giemsa solution (Sigma-Aldrich).

#### Flow cytometry analysis

Differentiated cells were gently pipetted and filtered through a 70 or 100 μm strainer sitting on a 50 mL tube. The cells were then pelleted by centrifugation and washed twice with PBS -/- solution containing 1% bovine serum albumin (BSA). The cells were stained with appropriate conjugated antibodies for 25 min at room temperature in dark, and analyzed in an Accuri C6 plus cytometer (Beckton Dickinson) after washing with BSA-containing PBS -/- solution. FlowJo software was used to process the collected flow data.

#### Transwell migration assay

Differentiated neutrophils were resuspended in HBSS buffer and allowed to migration for 2 hr towards fMLP (10 nM and 100 nM). Cells that migrated to the lower chamber were released with 0.5 M EDTA and counted using Accuri C6 plus cytometer (Beckton Dickinson). Live neutrophils were gated and analyzed in FlowJo software. The neutrophil counts were then normalized by the total numbers of cells added to each well.

#### 2D chemotaxis assay

Differentiated neutrophils were resuspended in HBBS with 20 mM HEPES and 0.5% FBS, and loaded into collagen-coated IBIDI chemotaxis μ-slides, which were then incubated at 37 °C for 30 min for cells to adhere. 15 μL of 1000 nM fMLP was loaded into the right reservoir yielding a final fMLP concentration of 187 nM. Cell migration was recorded every 60 s for a total of 120 min using LSM 710 (with Ziess EC Plan-NEOFLUAR 10X/0.3 objective) at 37 °C. Cells were tracked with ImageJ plug-in MTrackJ.

#### Phagocytosis

Phagocytosis was assessed using pHrodo Green *E.coli* BioParticles Conjugate according to the manufacturer’s protocol. In brief, pHrodo Green *E. coli* beads were resuspended in 2 mL of PBS and sonicated with an ultrasonicator 3 times. Beads per assay (100 mL) were opsonized by mixing with opsonizing reagent at a 1:1 ratio and incubated at 37 °C for 1 hr. Beads were washed 3 times with mHBSS buffer by centrifugation at 4 °C, 1,500 RCF for 15 min, and resuspended in mHBSS buffer. Differentiated neutrophils were resuspended in 100 μL of opsonized based solution and incubated at 37 °C for 1 hr, followed by flow cytometry analysis using a Accuri C6 plus cytometer (Beckton Dickinson).

#### Neutrophil-mediated *in vitro* cytotoxicity assay

The cell viability was analyzed by flow cytometry. In brief, 100 μL of tumor cells (50,000 cells/mL) were mixed with 100 μL of 150,000, 250,000 and 500,000 cells/mL neutrophils in 96 well plates, and then incubated at 37 °C, 5% CO2 for 24 hr. For the Cytochalasin D (Cayman Chemical) treatment, the neutrophils were firstly treated with 5 μM of Cytochalasin D before they incubated with tumor cells. For the N-acetylcysteine (NAC, Cayman Chemical) treatment, 5 mM of NAC were added during the incubation of neutrophils with tumor cells. For propofol treatment, 1, 3, 5, and 8 μg/mL of propofol was added during the incubation of neutrophils with tumor cells. To harvest all the cells, cell-containing media was firstly transferred into a new round-bottom 96-well plate, and 50 μL of trypsin-EDTA was added to the empty wells. After a 5-min incubation at 37 °C, attached cells were dissociated and transferred into the same wells of round-bottom 96-well plate with suspension cultures. All of the cells were pelleted by centrifuging the 96-well plate at 300 ×g, 4 °C for 4 min, and washed with 200 μL of PBS-/- solution containing 0.5% BSA. The pelleted cell mixtures were then stained with CD45 antibody and Calcein AM for 30 min at room temperature, and analyzed in the Accuri C6 plus cytometer (Beckton Dickinson).

#### Inducible gene knockdown in glioblastoma cells

To achieve inducible gene knockdown in glioblastoma cells, a PiggyBac (PB)-based inducible Cas13d plasmid (Addgene #155184) was implemented. The *CLCN3, ANXA2*, and *MMP2* targeting sgRNAs were designed using an online tool (https://cas13design.nygenome.org/) and cloned into the gRNA backbone to make *CLCN3, ANXA2*, and *MMP2* targeting plasmids (Addgene #170824-170830). The resulting sgRNA plasmids along with the hyPBase (kindly provided by Dr. Pentao Liu) and Cas13d plasmid were then introduced into U87MG cells via PEI transfection. After 2 to 4 days, transfected cells were treated with 5 μg/mL puromycin for one or two days to select drug-resistant tumor cells. After recovering, survived tumor cells were maintained under puromycin condition to avoid potential silencing of the integrated transgenes.

#### Bulk RNA sequencing and data analysis

Total RNA of sorted hPSC-derived CD16+ neutrophils was prepared with the Direct-zol RNA MiniPrep Plus kit (Zymo Research) according to the manufacturer’s instructions. RNA samples were then prepared and performed in Illumina HiSeq 2500 by the Center for Medical Genomics at Indiana University. HISAT2 program (Kim et al., 2019) was employed to map the resulting sequencing reads to the human genome (hg 19), and the python script rpkmforgenes.py (Ramsköld et al., 2009) was used to quantify the RefSeq transcript levels (RPKMs). The original fastq files and processed RPKM text files were submitted to NCBI GEO (GSE188393). RNA-seq data of human primary neutrophil samples (Perez et al., 2020) and hPSCs (Prasain et al., 2014; Tadeu et al., 2015) were retrieved previous studies. Heatmaps of selected gene subsets after normalization were then plotted using Morpheus (Broad Institute).

#### Conjugate formation assay

To visualize immunological synapses, 100 μL of U87MG cells (50,000 cells/mL) were seeded onto wells of 96-well plate and incubated at 37 °C for 12 hours, allowing them to attach. 100 μL neutrophils (500,000 cells/mL) were then added onto the target U87MG cells and incubated for 6 hours before fixation with 4% paraformaldehyde (in PBS). Cytoskeleton staining was then performed using an F-actin Visualization Biochem Kit (Cytoskeleton Inc.).

#### Trogocytosis assay

The transfer of membrane and cellular content from tumor cell to neutrophils was investigated using both microscope and flow cytometry analysis. Target tumor cells were labeled with Calcein-AM (1 μM) for 30 min at 37 °C. After washing with PBS, Calcein-AM labelled tumor cells were then incubated with neutrophils at a neutrophil-to-tumor ratio of 10:1. At different time points (between 0 to 6 hours), the resulting co-culture samples were imaged by a Leica DMi-8 fluorescent microscope. For the Cytochalasin D treatment, the neutrophils were pretreated with 5 μM of Cytochalasin D (Cayman Chemical) for 3 hours before they incubated with Calcein-AM labelled tumor cells. The floating neutrophils were collected for CD45 staining and analyzed in the Accuri C6 plus flow cytometer (Beckton Dickinson) after washing with PBS-/- solution containing 0.5% BSA.

#### Measurement of neutrophil extracellular trap (NET) formation

100 μL of U87MG cells (30,000 cells/mL) were seeded into wells of a 96-well plate 12 hours before adding neutrophils at a neutrophil-to-tumor ratio of 10:1. For propofol (Cayman Chemical) treatment, 1, 3, 5, and 8 μg/mL of propofol was added during the incubation of neutrophils with tumor cells. After co-incubation for 12 hours, the cells were centrifuged for 5 minutes at 200 Xg, and extracellular DNA in the supernatant samples was quantified using the Quant-iT PicoGreen™ dsDNA Assay Kit (Invitrogen) and characterized by SpectraMax iD3 microplate reader (Molecular Devices, Sunnyvale, CA, USA).

#### Measurement of reactive oxygen species (ROS) production in neutrophils

100 μL of U87MG cells (30,000 cells/mL) were seeded into wells of a 96-well plate 12 hours before adding neutrophils at a neutrophil-to-tumor ratio of 10:1. For the NAC treatment, 5 mM of NAC were added during the incubation of neutrophils with tumor cells. After co-incubation for 12 hours, the resulting cell mixture was treated with 10 μM H_2_DCFDA at 37 °C for 50 min and then the fluorescence emission signal (480-600 nm) was collected in a SpectraMax iD3 microplate reader (Molecular Devices, Sunnyvale, CA, USA) with an excitation wavelength of 475 nm.

#### Blood-brain-barrier (BBB) transmigration assay

*In vitro* BBB model was constructed with HBEC-5i cells in a transwell cell culture plate. Briefly, HBEC-5i cells (1×10^5^ cells/well) were seeded onto the upper chamber of the transwell pre-coated with gelatin (1% w:v) in 24-well transwell plates (8 μm pore size, 6.5 mm diameter, Corning), and maintained in DMEM/F12 medium containing 10% FBS. 2×10^5^ neutrophils were then added to the upper chamber, and FBS-free medium with or without fMLP (10 nM) was added to the lower chamber. After 3 hours of incubation, cell cultures were collected from the upper or lower chamber for the calculation of neutrophil numbers. For the cytotoxicity analysis, 2×10^4^ U87MG cells were seeded at the lower chamber 12 hours before adding neutrophils (2×10^5^ cells) to the upper chamber, and FBS-free medium with fMLP (10 nM) was then added to the lower chamber. After 12 hours of incubation, tumor cell viability was determined by flow cytometry analysis. For the second migration analysis, 2×10^5^ neutrophils from the bottom chamber of first migration were seeded on the upper chamber of second transwell BBB model, and the migrated neutrophils toward target tumor cells in the bottom chamber was quantified.

#### Neutrophil infiltration of 3D tumor spheroids

The 3D tumor spheroids were obtained using the hanging drop method. Briefly, U87MG cells were suspended in MEM medium with 10% FBS and 0.3% methylcellulose at 2×10^6^ cells/mL and deposited onto an inverted lid of 96-well plate as an individual drop using a 20 μL pipettor. The cover lid was then placed back onto the PBS-filled bottom chamber and incubated at 37 °C and 5% CO2. The hanging drops were monitored daily until cell aggregates were formed in ∼5-7 days. Each cell aggregate was transferred to a single well of a 24-well plate for the substream analysis. To assess the tumor penetration capability of CLTX T-CAR neutrophils, 2×10^5^ neutrophils/well were added to the wells of 24-well plate and incubated with the tumor spheroids. For the matrix metallopeptidase-2 (MMP2), the CLTX T-CAR neutrophils were treated with 50 ng/mL of MMP2 for 3 hours before they incubated with 3D tumor model. For the Cytochalasin D treatment, the CLTX T-CAR neutrophils were pretreated with 5 μM of Cytochalasin D for 3 hours before they incubated with 3D tumor cells. After co-incubation for 24 hours, the tumor spheroids were fixed and stained for CD45 and DAPI. For the cytotoxicity analysis, both live/dead staining and CytoTox-GloTM Cytotoxicity Assay kit (Promega) were employed. For the live/dead staining, the mixture of neutrophils and tumor spheroids were stained with 1 μM Calcein-AM (Invitrogen) and 1 μM propidium iodide (PI, Invitrogen). The stained cells were then imaged using a Leica DMi-8 fluorescent microscope. CytoTox-GloTM Cytotoxicity Assay was characterized by SpectraMax iD3 microplate reader (Molecular Devices, Sunnyvale, CA, USA).

### QUANTIFICATION AND STATISTICAL ANALYSIS

Data are presented as mean ± standard deviation (SD). Statistical significance was determined by Student’s t-test (two-tail) between two groups, and three or more groups were analyzed by one-way analysis of variance (ANOVA). P<0.05 was considered statistically significant.

## REFERENCE

1. Afonso, P. V., McCann, C.P., Kapnick, S.M., and Parent, C.A. (2013). Discoidin domain receptor 2 regulates neutrophil chemotaxis in 3D collagen matrices. Blood 121, 1644–1650.

2. Bao, X., Lian, X., Dunn, K.K., Shi, M., Han, T., Qian, T., Bhute, V.J., Canfield, S.G., and Palecek, S.P. (2015). Chemically-defined albumin-free differentiation of human pluripotent stem cells to endothelial progenitor cells. Stem Cell Res. 15, 122–129.

3. Bertrand, J.Y., Chi, N.C., Santoso, B., Teng, S., Stainier, D.Y.R., and Traver, D. (2010). Haematopoietic stem cells derive directly from aortic endothelium during development. Nature 464, 108–111.

4. Boisset, J.-C., van Cappellen, W., Andrieu-Soler, C., Galjart, N., Dzierzak, E., and Robin, C. (2010). In vivo imaging of haematopoietic cells emerging from the mouse aortic endothelium. Nature 464, 116–120.

5. Brok-Volchanskaya, V.S., Bennin, D.A., Suknuntha, K., Klemm, L.C., Huttenlocher, A., and Slukvin, I. (2019). Effective and Rapid Generation of Functional Neutrophils from Induced Pluripotent Stem Cells Using ETV2-Modified mRNA. Stem Cell Reports 13, 1099–1110.

6. Cao, X., Yakala, G.K., van den Hil, F.E., Cochrane, A., Mummery, C.L., and Orlova, V. V. (2019). Differentiation and Functional Comparison of Monocytes and Macrophages from hiPSCs with Peripheral Blood Derivatives. Stem Cell Reports 12, 1282–1297.

7. Chan, X.Y., Volkova, E., Eoh, J., Black, R., Fang, L., Gorashi, R., Song, J., Wang, J., Elliott, M.B., Barreto-Ortiz, S.F., et al. (2021). HIF2A gain-of-function mutation modulates the stiffness of smooth muscle cells and compromises vascular mechanics. IScience 24.

8. Chang, Y., Hellwarth, P.B., Randolph, L.N., Sun, Y., Xing, Y., Zhu, W., Lian, X.L., and Bao, X. (2020). Fluorescent indicators for continuous and lineage-specific reporting of cell-cycle phases in human pluripotent stem cells. Biotechnol. Bioeng. bit.27352.

9. Chen, Y., Cao, J., Xiong, M., Petersen, A.J., Dong, Y., Tao, Y., Huang, C.L.C.T.C.T.-L., Du, Z., Zhang, S.C.S.-C., Acampora, D., et al. (2015). Engineering Human Stem Cell Lines with Inducible Gene Knockout using CRISPR/Cas9. Cell Stem Cell 17, 233–244.

10. Choi, K.D., Vodyanik, M.A., and Slukvin, I.I. (2009). Generation of mature human myelomonocytic cells through expansion and differentiation of pluripotent stem cell-derived lin-CD34+CD43 +CD45+ progenitors. J. Clin. Invest. 119, 2818–2829.

11. Clarke, R.L., Yzaguirre, A.D., Yashiro-Ohtani, Y., Bondue, A., Blanpain, C., Pear, W.S., Speck, N.A., and Keller, G. (2013). The expression of Sox17 identifies and regulates haemogenic endothelium. Nat. Cell Biol. 15, 502–510.

12. Coffelt, S.B., Kersten, K., Doornebal, C.W., Weiden, J., Vrijland, K., Hau, C.S., Verstegen, N.J.M., Ciampricotti, M., Hawinkels, L.J.A.C., Jonkers, J., et al. (2015). IL-17-producing γδ T cells and neutrophils conspire to promote breast cancer metastasis. Nature 522, 345–348.

13. Coffelt, S.B., Wellenstein, M.D., and De Visser, K.E. (2016). Neutrophils in cancer: Neutral no more. Nat. Rev. Cancer 16, 431–446.

14. Devlin, J.C., Zwack, E.E., Tang, M.S., Li, Z., Fenyo, D., Torres, V.J., Ruggles, K. V., and Loke, P. (2020). Distinct Features of Human Myeloid Cell Cytokine Response Profiles Identify Neutrophil Activation by Cytokines as a Prognostic Feature during Tuberculosis and Cancer. J. Immunol. 204, 3389–3399.

15. Eruslanov, E.B., Singhal, S., and Albelda, S.M. (2017). Mouse versus Human Neutrophils in Cancer: A Major Knowledge Gap. Trends in Cancer 3, 149–160.

16. Esmann, L., Idel, C., Sarkar, A., Hellberg, L., Behnen, M., Möller, S., van Zandbergen, G., Klinger, M., Köhl, J., Bussmeyer, U., et al. (2010). Phagocytosis of Apoptotic Cells by Neutrophil Granulocytes: Diminished Proinflammatory Neutrophil Functions in the Presence of Apoptotic Cells. J. Immunol. 184, 391–400.

17. Feins, S., Kong, W., Williams, E.F., Milone, M.C., and Fraietta, J.A. (2019). An introduction to chimeric antigen receptor (CAR) T-cell immunotherapy for human cancer. Am. J. Hematol. 94, S3–S9.

18. Fidanza, A., Romanò, N., Ramachandran, P., Tamagno, S., Lopez-Yrigoyen, M., Taylor, A.H., Easterbrook, J., Henderson, B., Axton, R., Henderson, N.C., et al. (2019). Single cell transcriptome analysis reveals markers of naïve and lineage-primed hematopoietic progenitors derived from human pluripotent stem cells. BioRxiv 602565.

19. Fridlender, Z.G., Sun, J., Kim, S., Kapoor, V., Cheng, G., Ling, L., Worthen, G.S., and Albelda, S.M. (2009). Polarization of Tumor-Associated Neutrophil Phenotype by TGF-β: “N1” versus “N2” TAN. Cancer Cell 16, 183–194.

20. Hayashi, F., Means, T.K., and Luster, A.D. (2003). Toll-like receptors stimulate human neutrophil function. Blood 102, 2660–2669.

21. Huang, X., Trinh, T., Aljoufi, A., and Broxmeyer, H.E. (2018). Hypoxia Signaling Pathway in Stem Cell Regulation: Good and Evil. Curr. Stem Cell Reports 4, 149–157.

22. Huo, X., Li, H., Li, Z., Yan, C., Agrawal, I., Mathavan, S., Liu, J., and Gong, Z. (2019). Transcriptomic profiles of tumor-associated neutrophils reveal prominent roles in enhancing angiogenesis in liver tumorigenesis in zebrafish. Sci. Rep. 9.

23. Ilie, M., Hofman, V., Ortholan, C., Bonnetaud, C., Coëlle, C., Mouroux, J., and Hofman, P. (2012). Predictive clinical outcome of the intratumoral CD66b-positive neutrophil-to-CD8-positive T-cell ratio in patients with resectable nonsmall cell lung cancer. Cancer 118, 1726–1737.

24. Itatani, Y., Yamamoto, T., Zhong, C., Molinolo, A.A., Ruppel, J., Hegde, P., Taketo, M.M., and Ferrara, N. (2020). Suppressing neutrophil-dependent angiogenesis abrogates resistance to anti-VEGF antibody in a genetic model of colorectal cancer. Proc. Natl. Acad. Sci. U. S. A. 117, 21598–21608.

25. Itoh, Y. (2015). Membrane-type matrix metalloproteinases: Their functions and regulations. Matrix Biol.

26. Jaillon, S., Ponzetta, A., Di Mitri, D., Santoni, A., Bonecchi, R., and Mantovani, A. (2020). Neutrophil diversity and plasticity in tumour progression and therapy. Nat. Rev. Cancer 20, 485–503.

27. June, C.H., and Sadelain, M. (2018). Chimeric Antigen Receptor Therapy. N. Engl. J. Med. 379, 64–73.

28. June, C.H., O’Connor, R.S., Kawalekar, O.U., Ghassemi, S., and Milone, M.C. (2018). CAR T cell immunotherapy for human cancer. Science (80-.). 359, 1361–1365.

29. Kalafati, L., Kourtzelis, I., Schulte-Schrepping, J., Li, X., Hatzioannou, A., Grinenko, T., Hagag, E., Sinha, A., Has, C., Dietz, S., et al. (2020). Innate Immune Training of Granulopoiesis Promotes Anti-tumor Activity. Cell 183, 771–785.e12.

30. Kargl, J., Zhu, X., Zhang, H., Yang, G.H.Y., Friesen, T.J., Shipley, M., Maeda, D.Y., Zebala, J.A., McKay-Fleisch, J., Meredith, G., et al. (2019). Neutrophil content predicts lymphocyte depletion and anti-PD1 treatment failure in NSCLC. JCI Insight 4.

31. Kim, D., Paggi, J.M., Park, C., Bennett, C., and Salzberg, S.L. (2019). Graph-based genome alignment and genotyping with HISAT2 and HISAT-genotype. Nat. Biotechnol. 37, 907–915.

32. Kim, G.B., Aragon-Sanabria, V., Randolph, L., Jiang, H., Reynolds, J.A., Webb, B.S., Madhankumar, A., Lian, X., Connor, J.R., Yang, J., et al. (2020a). High-affinity mutant Interleukin-13 targeted CAR T cells enhance delivery of clickable biodegradable fluorescent nanoparticles to glioblastoma. Bioact. Mater. 5, 624–635.

33. Kim, G.B., Aragon-Sanabria, V., Randolph, L., Jiang, H., Reynolds, J.A., Webb, B.S., Madhankumar, A., Lian, X., Connor, J.R., Yang, J., et al. (2020b). High-affinity mutant Interleukin-13 targeted CAR T cells enhance delivery of clickable biodegradable fluorescent nanoparticles to glioblastoma. Bioact. Mater. 5, 624–635.

34. Kim, I., Saunders, T.L., and Morrison, S.J. (2007). Sox17 Dependence Distinguishes the Transcriptional Regulation of Fetal from Adult Hematopoietic Stem Cells. Cell 130, 470–483.

35. Kissa, K., and Herbomel, P. (2010). Blood stem cells emerge from aortic endothelium by a novel type of cell transition. Nature 464, 112–115.

36. Klichinsky, M., Ruella, M., Shestova, O., Lu, X.M., Best, A., Zeeman, M., Schmierer, M., Gabrusiewicz, K., Anderson, N.R., Petty, N.E., et al. (2020). Human chimeric antigen receptor macrophages for cancer immunotherapy. Nat. Biotechnol. 1–7.

37. Lachmann, N., Ackermann, M., Frenzel, E., Liebhaber, S., Brennig, S., Happle, C., Hoffmann, D., Klimenkova, O., Lüttge, D., Buchegger, T., et al. (2015). Large-scale hematopoietic differentiation of human induced pluripotent stem cells provides granulocytes or macrophages for cell replacement therapies. Stem Cell Reports 4, 282–296.

38. Lawrence, S.M., Corriden, R., and Nizet, V. (2018). The Ontogeny of a Neutrophil: Mechanisms of Granulopoiesis and Homeostasis. Microbiol. Mol. Biol. Rev. 82.

39. Lecot, P., Sarabi, M., Pereira Abrantes, M., Mussard, J., Koenderman, L., Caux, C., Bendriss-Vermare, N., and Michallet, M.C. (2019). Neutrophil Heterogeneity in Cancer: From Biology to Therapies. Front. Immunol. 10.

40. Li, L., Qi, X., Sun, W., Abdel-Azim, H., Lou, S., Zhu, H., Prasadarao, N. V, Zhou, A., Shimada, H., Shudo, K., et al. (2016). Am80-GCSF synergizes myeloid expansion and differentiation to generate functional neutrophils that reduce neutropenia-associated infection and mortality. EMBO Mol. Med. 8, 1340–1359.

41. Li, Y., Hermanson, D.L., Moriarity, B.S., and Kaufman, D.S. (2018). Human iPSC-Derived Natural Killer Cells Engineered with Chimeric Antigen Receptors Enhance Anti-tumor Activity. Cell Stem Cell 23, 181–192.e5.

42. Lian, X., Zhang, J., Azarin, S.M., Zhu, K., Hazeltine, L.B., Bao, X., Hsiao, C., Kamp, T.J., and Palecek, S.P. (2013). Directed cardiomyocyte differentiation from human pluripotent stem cells by modulating Wnt/β-catenin signaling under fully defined conditions. Nat. Protoc. 8, 162–175.

43. Lian, X., Bao, X., Al-Ahmad, A., Liu, J., Wu, Y., Dong, W., Dunn, K.K., Shusta, E.V., and Palecek, S.P. (2014). Efficient Differentiation of Human Pluripotent Stem Cells to Endothelial Progenitors via Small-Molecule Activation of WNT Signaling. Stem Cell Reports 3, 804–816.

44. Liang, J., Piao, Y., Holmes, L., Fuller, G.N., Henry, V., Tiao, N., and De Groot, J.F. (2014). Neutrophils promote the malignant glioma phenotype through S100A4. Clin. Cancer Res. 20, 187–198.

45. Lim, W.A., and June, C.H. (2017). The Principles of Engineering Immune Cells to Treat Cancer. Cell 168, 724–740.

46. Lyons, S.A., O’Neal, J., and Sontheimer, H. (2002). Chlorotoxin, a scorpion-derived peptide, specifically binds to gliomas and tumors of neuroectodermal origin. Glia 39, 162–173.

47. Matlung, H.L., Babes, L., Zhao, X.W., van Houdt, M., Treffers, L.W., van Rees, D.J., Franke, K., Schornagel, K., Verkuijlen, P., Janssen, H., et al. (2018). Neutrophils Kill Antibody-Opsonized Cancer Cells by Trogoptosis. Cell Rep. 23, 3946–3959.e6.

48. McDermott, D.H., De Ravin, S.S., Jun, H.S., Liu, Q., Long Priel, D.A., Noel, P., Takemoto, C.M., Ojode, T., Paul, S.M., Dunsmore, K.P., et al. (2010). Severe congenital neutropenia resulting from G6PC3 deficiency with increased neutrophil CXCR4 expression and myelokathexis. Blood 116, 2793–2802.

49. Mehta, R.S., and Rezvani, K. (2018). Chimeric antigen receptor expressing natural killer cells for the immunotherapy of cancer. Front. Immunol. 9.

50. Meier, A., Chien, J., Hobohm, L., Patras, K.A., Nizet, V., and Corriden, R. (2019). Inhibition of human neutrophil extracellular trap (NET) production by propofol and lipid emulsion. Front. Pharmacol. 10, 323.

51. Neubert, E., Meyer, D., Rocca, F., Günay, G., Kwaczala-Tessmann, A., Grandke, J., Senger-Sander, S., Geisler, C., Egner, A., Schön, M.P., et al. (2018). Chromatin swelling drives neutrophil extracellular trap release. Nat. Commun. 9.

52. Ng, E.S., Azzola, L., Bruveris, F.F., Calvanese, V., Phipson, B., Vlahos, K., Hirst, C., Jokubaitis, V.J., Yu, Q.C., Maksimovic, J., et al. (2016). Differentiation of human embryonic stem cells to HOXA+ hemogenic vasculature that resembles the aorta-gonad-mesonephros. Nat. Biotechnol. 34, 1168–1179.

53. Nguyen, V., Conyers, J.M., Zhu, D., Gibo, D.M., Hantgan, R.R., Larson, S.M., Debinski, W., and Mintz, A. (2012). A novel ligand delivery system to non-invasively visualize and therapeutically exploit the IL13Rα2 tumor-restricted biomarker. Neuro. Oncol. 14, 1239–1253.

54. Oatley, M., Bölükbası, Ö.V., Svensson, V., Shvartsman, M., Ganter, K., Zirngibl, K., Pavlovich, P. V., Milchevskaya, V., Foteva, V., Natarajan, K.N., et al. (2020). Single-cell transcriptomics identifies CD44 as a marker and regulator of endothelial to haematopoietic transition. Nat. Commun. 11.

55. Perez, C., Botta, C., Zabaleta, A., Puig, N., Cedena, M.T., Goicoechea, I., Alameda, D., José-Eneriz, E.S., Merino, J., Rodríguez-Otero, P., et al. (2020). Immunogenomic identification and characterization of granulocytic myeloid-derived suppressor cells in multiple myeloma. Blood 136, 199–209.

56. Ponzetta, A., Carriero, R., Carnevale, S., Barbagallo, M., Molgora, M., Perucchini, C., Magrini, E., Gianni, F., Kunderfranco, P., Polentarutti, N., et al. (2019). Neutrophils Driving Unconventional T Cells Mediate Resistance against Murine Sarcomas and Selected Human Tumors. Cell 178, 346–360.e24.

57. Prasain, N., Lee, M.R., Vemula, S., Meador, J.L., Yoshimoto, M., Ferkowicz, M.J., Fett, A., Gupta, M., Rapp, B.M., Saadatzadeh, M.R., et al. (2014). Differentiation of human pluripotent stem cells to cells similar to cord-blood endothelial colony-forming cells. Nat. Biotechnol. 32, 1151–1157.

58. Qin, C., He, B., Dai, W., Zhang, H., Wang, X., Wang, J., Zhang, X., Wang, G., Yin, L., and Zhang, Q. (2014). Inhibition of metastatic tumor growth and metastasis via targeting metastatic breast cancer by chlorotoxin-modified liposomes. Mol. Pharm.

59. Ramsköld, D., Wang, E.T., Burge, C.B., and Sandberg, R. (2009). An abundance of ubiquitously expressed genes revealed by tissue transcriptome sequence data. PLoS Comput. Biol. 5, e1000598.

60. Rankin, E.B., Nam, J.M., and Giaccia, A.J. (2016). Hypoxia: Signaling the Metastatic Cascade. Trends in Cancer 2, 295–304.

61. Rincón, E., Rocha-Gregg, B.L., and Collins, S.R. (2018). A map of gene expression in neutrophil-like cell lines. BMC Genomics 2018 191 19, 1–17.

62. Roberts, M.R., Cooke, K.S., Tran, A.C., Smith, K.A., Lin, W.Y., Wang, M., Dull, T.J., Farson, D., Zsebo, K.M., and Finer, M.H. (1998). Antigen-specific cytolysis by neutrophils and NK cells expressing chimeric immune receptors bearing zeta or gamma signaling domains. J. Immunol. 161, 375–384.

63. Rosen, J., Blau, T., Grau, S.J., Barbe, M.T., Fink, G.R., and Galldiks, N. (2018). Extracranial Metastases of a Cerebral Glioblastoma: A Case Report and Review of the Literature. Case Rep. Oncol. 11, 591–600.

64. Saeki, K., Saeki, K., Nakahara, M., Matsuyama, S., Nakamura, N., Yogiashi, Y., Yoneda, A., Koyanagi, M., Kondo, Y., and Yuo, A. (2009). A Feeder-Free and Efficient Production of Functional Neutrophils from Human Embryonic Stem Cells. Stem Cells 27, 59–67.

65. Sagiv, J.Y., Michaeli, J., Assi, S., Mishalian, I., Kisos, H., Levy, L., Damti, P., Lumbroso, D., Polyansky, L., Sionov, R. V., et al. (2015). Phenotypic diversity and plasticity in circulating neutrophil subpopulations in cancer. Cell Rep. 10, 562–573.

66. Seo, Y.J., Cho, W.H., Kang, D.W., and Cha, S.H. (2012). Extraneural metastasis of glioblastoma multiforme presenting as an unusual neck mass. J. Korean Neurosurg. Soc. 51, 147–150.

67. Shaul, M.E., Levy, L., Sun, J., Mishalian, I., Singhal, S., Kapoor, V., Horng, W., Fridlender, G., Albelda, S.M., and Fridlender, Z.G. (2016). Tumor-associated neutrophils display a distinct N1 profile following TGFβ modulation: A transcriptomics analysis of pro-vs. antitumor TANs. Oncoimmunology 5.

68. Smith, C., Gore, A., Yan, W., Abalde-Atristain, L., Li, Z., He, C., Wang, Y., Brodsky, R.A., Zhang, K., Cheng, L., et al. (2014). Whole-Genome Sequencing Analysis Reveals High Specificity of CRISPR/Cas9 and TALEN-Based Genome Editing in Human iPSCs. Cell Stem Cell 15, 12–13.

69. Smith, J.R., Maguire, S., Davis, L.A., Alexander, M., Yang, F., Chandran, S., ffrench-Constant, C., and Pedersen, R.A. (2008). Robust, Persistent Transgene Expression in Human Embryonic Stem Cells Is Achieved with AAVS1-Targeted Integration. Stem Cells 26, 496–504.

70. Sweeney, C.L., Teng, R., Wang, H., Merling, R.K., Lee, J., Choi, U., Koontz, S., Wright, D.G., and Malech, H.L. (2016). Molecular Analysis of Neutrophil Differentiation from Human Induced Pluripotent Stem Cells Delineates the Kinetics of Key Regulators of Hematopoiesis. Stem Cells 34, 1513–1526.

71. Tadeu, A.M.B., Lin, S., Hou, L., Chung, L., Zhong, M., Zhao, H., and Horsley, V. (2015). Transcriptional profiling of ectoderm specification to keratinocyte fate in human embryonic stem cells. PLoS One 10, e0122493.

72. Trump, L.R., Nayak, R.C., Singh, A.K., Emberesh, S., Wellendorf, A.M., Lutzko, C.M., and Cancelas, J.A. (2019). Neutrophils Derived from Genetically Modified Human Induced Pluripotent Stem Cells Circulate and Phagocytose Bacteria In Vivo. Stem Cells Transl. Med. 8, 557–567.

73. Veres, A., Gosis, B.S., Ding, Q., Collins, R., Ragavendran, A., Brand, H., Erdin, S., Talkowski, M.E., and Musunuru, K. (2014). Low Incidence of Off-Target Mutations in Individual CRISPR-Cas9 and TALEN Targeted Human Stem Cell Clones Detected by Whole-Genome Sequencing. Cell Stem Cell 15, 27–30.

74. Wang, D., Starr, R., Chang, W.C., Aguilar, B., Alizadeh, D., Wright, S.L., Yang, X., Brito, A., Sarkissian, A., Ostberg, J.R., et al. (2020a). Chlorotoxin-directed CAR T cells for specific and effective targeting of glioblastoma. Sci. Transl. Med. 12.

75. Wang, J., Toregrosa-Allen, S., Elzey, B.D., Utturkar, S., Lanman, N.A., Bernal-Crespo, V., Behymer, M.M., Knipp, G.T., Yun, Y., Veronesi, M.C., et al. (2020b). Tumor-responsive, multifunctional CAR-NK cells cooperate with impaired autophagy to infiltrate and target glioblastoma. BioRxiv 2020.10.07.330043.

76. Wang, T., Cao, L., Dong, X., Wu, F., De, W., Huang, L., and Wan, Q. (2020c). LINC01116 promotes tumor proliferation and neutrophil recruitment via DDX5-mediated regulation of IL-1β in glioma cell. Cell Death Dis. 11.

77. Wu, M., Zhang, H., Tie, C., Yan, C., Deng, Z., Wan, Q., Liu, X., Yan, F., and Zheng, H. (2018). MR imaging tracking of inflammation-activatable engineered neutrophils for targeted therapy of surgically treated glioma. Nat. Commun. 9, 1–13.

78. Xie, X., Shi, Q., Wu, P., Zhang, X., Kambara, H., Su, J., Yu, H., Park, S.Y., Guo, R., Ren, Q., et al. (2020). Single-cell transcriptome profiling reveals neutrophil heterogeneity in homeostasis and infection. Nat. Immunol. 21, 1119–1133.

79. Xue, J., Zhao, Z., Zhang, L., Xue, L., Shen, S., Wen, Y., Wei, Z., Wang, L., Kong, L., Sun, H., et al. (2017). Neutrophil-mediated anticancer drug delivery for suppression of postoperative malignant glioma recurrence. Nat. Nanotechnol. 12, 692–700.

80. Yan, J., Kloecker, G., Fleming, C., Bousamra, M., Hansen, R., Hu, X., Ding, C., Cai, Y., Xiang, D., Donninger, H., et al. (2014). Human polymorphonuclear neutrophils specifically recognize and kill cancerous cells. Oncoimmunology 3, e950163.

81. Zhao, Y., Rahmy, S., Liu, Z., Zhang, C., and Lu, X. (2020). Rational targeting of immunosuppressive neutrophils in cancer. Pharmacol. Ther. 212.

82. Zhu, H., Lai, Y.-S., Li, Y., Blum, R.H., and Kaufman, D.S. (2018). Concise Review: Human Pluripotent Stem Cells to Produce Cell-Based Cancer Immunotherapy. Stem Cells 36, 134–145.

